# CRISPR screens of noncanonical coding sequences and integrative multiomics uncover novel retrovirus-derived proteins

**DOI:** 10.64898/2026.02.23.707583

**Authors:** Eric Malekos, Valeriya Smaliy, Lisa Sudek, Susan Carpenter

## Abstract

Ribosome profiling has revealed thousands of noncanonical translation events across mammalian genomes, yet functional characterization has overwhelmingly focused on proliferative fitness in cancer cell lines. Here, we present a comprehensive survey of noncanonical translation in the mouse immune system and its functional consequences in macrophages. By performing a meta-analysis of Ribo-seq across 20 public mouse leukocyte and 5 human leukocyte datasets we define a compendium of 22,572 mouse and 18,701 human noncanonical coding sequences (nCDSs), including upstream ORFs (uORFs), downstream ORFs, and ORFs on noncoding RNAs and pseudogenes (ncORFs). Cross species conservation and proteogenomic integration with reanalyzed mass spectrometry data prioritizes a high-confidence subset. To move beyond cataloging, we carried out two orthogonal CRISPR screens in immortalized bone marrow-derived macrophages: a fitness screen identifying nCDSs required for macrophage viability, and a TLR1/TLR2-NFκB reporter screen uncovering nCDSs that modulate innate immune signaling. These screens suggested dozens of nCDSs for functional follow up including a family of endogenous retroviral envelope-derived proteins translated in adult mouse myeloid cells. Updated single-cell RNA-seq annotations and an interactive UCSC Genome Browser session integrating Ribo-seq, proteomics, and CRISPR screen data are provided as community resources. Together, these findings expand the functional landscape of noncanonical translation in immunity and nominate endogenous retroviral proteins as previously unrecognized regulators of murine macrophage biology.

## Introduction

Over the past decade, ribosome profiling (Ribo-seq) has revealed that translation occurs at thousands of sites beyond canonical coding sequences. In mRNAs this includes translation of upstream open reading frames (uORFs) in the 5’ untranslated region (UTR), internal ORFs (iORFs) nested in the coding sequence (CDS), and downstream ORFs (dORFs) in the 3’ UTR. Beyond annotated coding genes, translation of ORFs embedded at loci annotated as pseudogenes and noncoding RNAs (ncORFs) has also been repeatedly documented^1–4^. In this report we will refer to the subset of ORFs which are predicted to be translated as noncanonical coding sequences (nCDSs). These nCDSs give rise to a “dark proteome”^5^ whose scope continues to expand with improved detection methods and deeper sequencing coverage, raising fundamental questions about which translated sequences carry biological function.

Early genome annotation efforts relied heavily on sequence length and evolutionary conservation, establishing a 100-codon threshold that systematically excluded short ORFs from reference gene sets^6^. As a consequence, there may be thousands of unannotated small proteins and regulatory peptides, arising from misclassified “noncoding” transcripts. Computational rescoring of genomic sequences, combined with advances in Ribo-seq and mass spectrometry-based proteomics, has since identified hundreds of translated nCDSs with important phenotypic effects, including defined roles in inflammation^7–9^, skeletal muscle activity^10,11^, and cell differentiation^12^. These discoveries have motivated efforts to catalog noncanonical translation events and to integrate them into updated genome annotations^13,14^.

Despite progress in cataloging nCDSs^2,15–17^, functional characterization has lagged substantially behind discovery. The vast majority of Ribo-seq-detected translation events remain uncharacterized with respect to biological activity. Moreover, many small, nCDS translation products are likely to be unstable, degrading shortly after production, and not act as functional effectors^14,18,19^. Efforts have been made to ascribe biological function to some of the tens of thousands of predicted nCDSs with a handful of high-throughput perturbation studies linking these ORFs to phenotypes. However, these screens have predominantly tested nCDS essentiality in proliferating cancer cell lines and stem cells^20–24^, probing nCDS importance in a relatively narrow context. A notable exception is a targeted Perturb-seq experiment in K562s which provided a rich readout of altered gene programs following nCDS knockout, albeit at reduced throughput compared to pooled screens^22^. Thus, while the laudable goal of identifying cancer driving noncanonical translation events has helped contextualize some of these nCDSs, the field has not witnessed similar efforts to apply high throughput screens in other biological contexts.

Immune cells represent a particularly compelling context in which to explore noncanonical translation^25^. Leukocytes must rapidly adapt their proteomes in response to inflammatory cues, pathogen encounter, and tissue damage. This plasticity is regulated in part at the translational level, with global reprogramming of ribosome occupancy occurring within hours of immune activation^26,27^. uORFs have emerged as key regulatory elements in this context, modulating translation of downstream CDSs for cytokines, transcription factors, and stress-response genes^28–34^. Moreover, long noncoding RNAs and pseudogenes can be induced in inflammatory contexts^35,36^, and the translational potential of these loci remains largely unexplored.

A related underexplored dimension of noncanonical translation concerns endogenous retroviruses (ERVs), remnants of ancient retroviral integrations that comprise roughly 8-10% of mammalian genomes^37,38^. Most ERV sequences have degenerated beyond coding capacity, but a subset retains intact ORFs and can be transcribed and translated^39^. The best-characterized examples are the syncytins, envelope-derived (Env) genes that have been co-opted for placental development and trophoblast fusion across placental mammalian lineages^40,41^. Whether additional ERV ORFs are translated outside the placenta, and how they might contribute to immune function, remain open questions.

Addressing these gaps required an integrative approach that combines detection of translation events and high-throughput functional assays. Here, we present a unified Ribo-seq meta-analysis of 20 public mouse leukocyte datasets available at the outset of this study to define a comprehensive catalog of nCDSs translated in the immune system. We merge computational translation predictions from two complementary callers with a custom transcriptome annotation and highlight hits with confident proteomics support. The pipeline is further applied to 5 Ribo-seq datasets in human leukocytes and leukocyte derived cell lines. We then carry out two orthogonal CRISPR screens in immortalized bone marrow derived macrophages (iBMDMs) - an essentiality screen for fitness effects and a TLR1/TLR2-NFκB reporter screen for inflammatory signaling - to identify nCDSs with measurable biological activity. This strategy nominates translated uORFs and ncORFs that modulate macrophage viability and innate immune activation, including several conserved between mouse and human. Finally, we characterize a family of translated Env proteins expressed in myeloid cells, including a syncytin-like membrane protein that tunes NFκB-responsive transcription and a secreted Env fragment with structural homology to the feline leukemia virus accessory (FeLIX) protein. Together, we present a first pass at annotating the functional landscape of noncanonical translation in immunity and establish resources, including updated single-cell annotations and an interactive genome browser, for continued exploration of noncanonical translation.

## Results

### A unified leukocyte Ribo-seq meta-analysis defines a noncanonical ORF compendium

We analyzed 20 publicly available Ribo-seq datasets spanning mouse leukocyte lineages (neutrophils^42^, macrophages^7,43–46^, B cells^47–50^, CD4^50–55^ and CD8 T cells^56^, and dendritic cells^57–59^) and kidney tissue^60^ treated with the inflammatory ligand lipopolysaccharide (LPS) (Fig. 1A,B). After preprocessing to remove contaminating reads (Supp. Fig. 1A,C, Methods), the majority of retained reads mapped in-frame to annotated coding sequence (CDS) p-sites, supporting expected ribosome occupancy (Supp. Fig. 1B,D).

**Figure 1.**
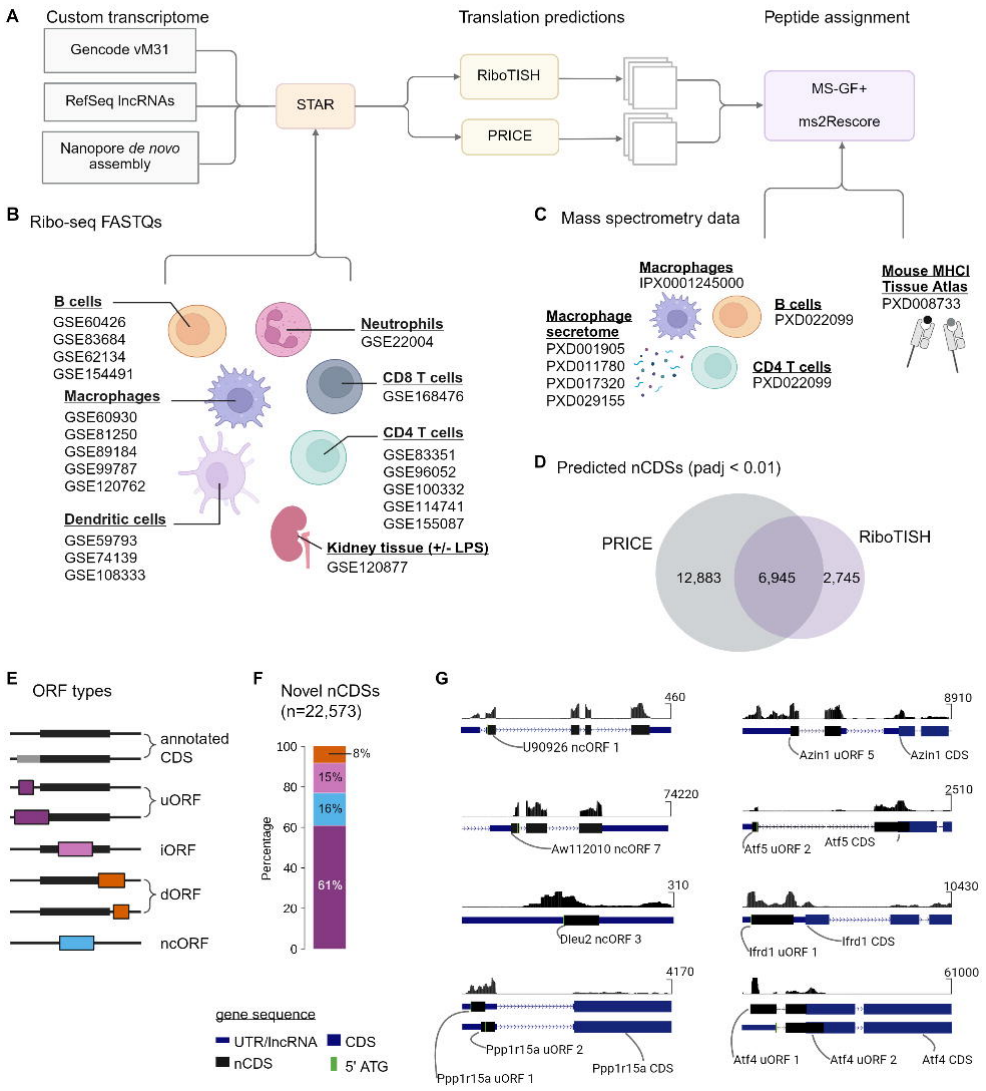
**Integrated Ribo-seq and proteomics pipeline across immune cells**. (A) Overview of the noncanonical CDS (nCDS) discovery pipeline. Ribo-seq reads were aligned with STAR to a custom transcriptome constructed from GENCODE vM31, RefSeq lncRNAs, and a Nanopore *de novo* assembly. Translation events were inferred independently using RiboTISH and PRICE. The predicted nCDS peptides were merged with GENCODE protein annotations to generate a custom spectral search database for MS-GF+ and ms2Rescore analysis of publicly available proteomics datasets. (B) Publicly available Ribo-seq datasets analyzed in this study. (C) Publicly available mass spectrometry datasets. (D) Shared predictions between RiboTISH- and PRICE-derived nCDSs at each tool’s Benjamini Hochberg adjusted p-value < 0.01 cut-off. (E) Representative nCDS categories identified by the pipeline. Canonical CDS entries and their N-terminal extensions were not considered in subsequent analysis. (F) Composition of the aggregated nCDS catalog across all datasets. (G) Reidentification of previously documented nCDSs. CDS: coding sequence; UTR: untranslated region; lncRNA: long noncoding RNA; uORF: upstream open reading frame; iORF: internal open reading frame; dORF: downstream open reading frame; ncORF: noncoding RNA open reading frame

We generated a custom transcriptome that merges GENCODE vM31, RefSeq lncRNAs, and a *de novo* Nanopore direct RNA assembly^61^, to enable comprehensive translation discovery (Fig. 1A). Translation events were inferred independently using two complementary callers - RiboTISH^62^ and PRICE^63^ (Fig. 1A; Tables S1,S2) - and predicted protein sequences were merged with GENCODE proteins to generate a tailored spectral search database for peptide spectra matching (Fig. 1A,C, Methods). Aggregating predictions across all datasets yielded a catalog of 22,572 nCDSs spanning uORFs, iORFs, dORFs, and ncORFs, while excluding annotated CDS and N-terminal extensions from the noncanonical set (Fig. 1E,F, Table S3).

UORFs made up the largest fraction of nCDS calls, consistent with previous reports^20,22,64^ and unsurprising given their position upstream of canonical CDSs and preferential access to the pre-initiation complex. At matched confidence thresholds (adjusted p-value [padj] < 0.01), PRICE proved more sensitive, identifying a larger number of ORFs and reidentifying 10 out of 10 previously identified nCDSs compared to 8 out of 10 reindentifications with RiboTISH (Fig. 1D,G, Table S4). We separately analyzed 5 public human Ribo-seq datasets: B cell derived^65,66^ (B721.221, Mutu), monocyte derived^67^ (THP-1), and CD8 T-cells^68,69^ in a pipeline that matched the mouse case (Table S5,S6; Methods).

Consistent with expectations for noncanonical translation, predicted nCDSs were predominantly short, with many below the historical 100-codon cut-off used in earlier coding sequence annotation^6^ (Supp. Fig. 2A,B). In silico localization predictions with DeepLoc^70^ further suggested that proteins encoded by predicted nCDSs differ from canonical proteomes, including an increased propensity for mitochondrial localization and reduced propensity for membrane embedding, consistent with prior nCDSs predictions^19^(Supp. Fig. 2C,D, Tables S8-S13).

**Figure 2.**
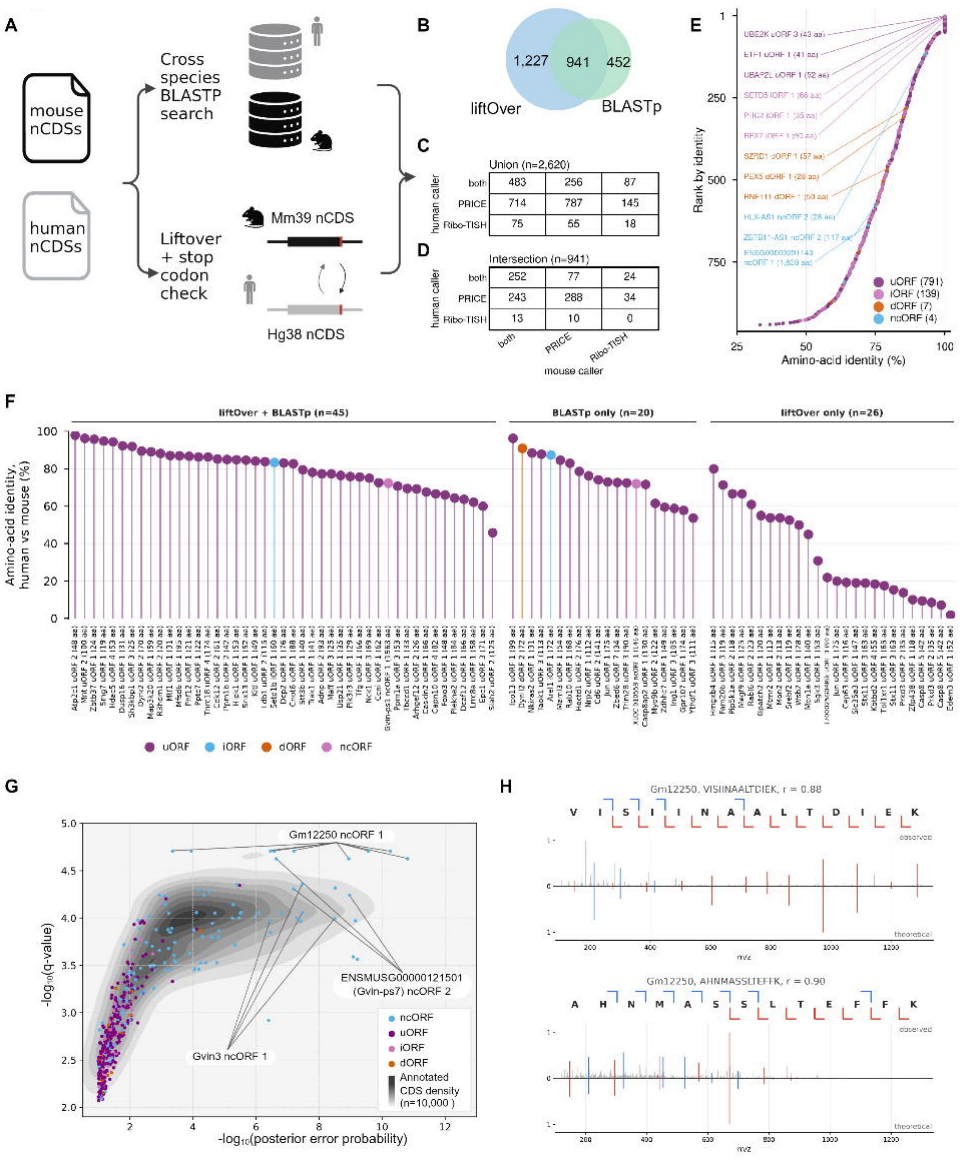
**Cross species conservations and proteomics supports translation of select noncanonical loci**. (A) NCDS conservation was tested by cross species BLASTp search and stop coordinate liftover mapping between species. BLASTp reciprocal best hits are reported as conserved when search in both directions returns the same nCDS and each E < 10^-5^. Liftover reciprocal best hits are reported when the nCDS stop codons map bidirectionally between species. (B) Venn diagram show concordance of conservation approaches (C,D) Tables depicting the (C) union of 2,620 nCDSs and (D) intersection of 941 nCDSs along with each nCDS caller. (E) The 941 conserved intersection hits ranked by amino acid identity. (F) Lollipop plot for nCDSs with both conservation and proteomics evidence, requiring a unique PSM with q-value < 0.01 and posterior error probability < 0.05 (G) PSMs of unique nCDSs found in macrophage cell lysate mass spectrometry dataset IPX0001245000 which includes macrophages from many tissues. The density cloud represents 10,000 randomly selected PSMs that mapped to protein coding genes. All displayed PSMs are significant at q-value < 0.01, posterior error probability < 0.1 cut-offs. (H) Peptide spectra that map uniquely to Gm12250, nowhere else in the genome, passed FDR filters, and scored “excellent” according to the heuristic scorer (Methods).

### Thousands of upstream ORFs are conserved between human and mouse

To determine whether noncanonical translation events are conserved, we compared the mouse compendium with nCDSs called from the human datasets using the same workflow. Human calls were first filtered against GENCODE v49 to remove ORFs that are in frame with an annotated CDS or share a start or stop codon with an annotated CDS, yielding 18,701 human nCDSs (Methods). We then paired human and mouse nCDSs along two orthogonal axes (Fig. 2A). For sequence homology, each nCDS peptide was searched with BLASTp^71^ against all nCDS peptides and the GENCODE proteome of the other species, and reciprocal best hits between two nCDSs (E < 10 in each direction) were retained; searches against shuffled decoy peptides estimated a false discovery rate (FDR) of <1% at this threshold. For positional homology, stop codons were projected between the mouse and human genomes with liftOver^72,73^ in both directions, and pairs whose stop codons mapped reciprocally were retained.

Together, these approaches identified 2,485 human-mouse nCDS pairs, 857 of which were supported by both sequence and position (Fig. 2B-D, Table S18). Conservation was dominated by uORFs: 2,051 human uORFs (19.7%) had a mouse counterpart, 715 of them by both criteria, compared with 15.4% of iORFs, 2.9% of dORFs and 0.6% of ncORFs. Pairs supported by both criteria shared a median of 79% amino-acid identity (Fig. 2E). Conserved nCDSs were also enriched for independent evidence of translation. In both species they were more often called by both RiboTISH and PRICE (human 39% versus 22%; mouse 54% versus 30%). Mouse nCDSs with peptide support (q-value < 0.01; PEP < 0.1) were conserved nearly twice as often as those without (20.7% versus 10.2%; Mantel-Haenszel odds ratio 1.77 after stratifying by ORF type and length, P = 7.2 × 10^-9^). Conserved ncORFs were rare but included ORFs on lncRNAs antisense to ZBTB11 and HLX (75% and 93% identity). Notably, the ncORF on the human gene ZBTB11-AS1 was previously demonstrated to have an essentiality function in cancer cell lines. They also included a 1,839-codon ORF in the human GVIN locus that aligned to the Gvin-ps1 ncORF (72% identity), consistent with translation of the interferon-inducible GVIN pseudogene family described below.

### Proteogenomic evidence strongly supports translation of pseudogenes and lncRNA-encoded zinc finger proteins

To determine which Ribo-seq-inferred translation events yield detectable protein products, we reanalyzed mouse mass spectrometry datasets with our predicted nCDS peptide database using MS-GF+^74^ for initial search and MS2Rescore^75^ to recalculate assignments (Fig. 1A; Tables S19,S20). High confidence peptide spectrum matches (PSMs) were reported (q-value < 0.01; posterior error probability < 0.1) and we highlighted nCDSs supported by either robust total peptide-spectrum matches (PSMs; ≥5) or multiple unique PSMs (≥2) and visualized their transcription across immune lineages using uniformly processed gene expression estimates (Supp. Fig. 3A; Table S22).

**Figure 3.**
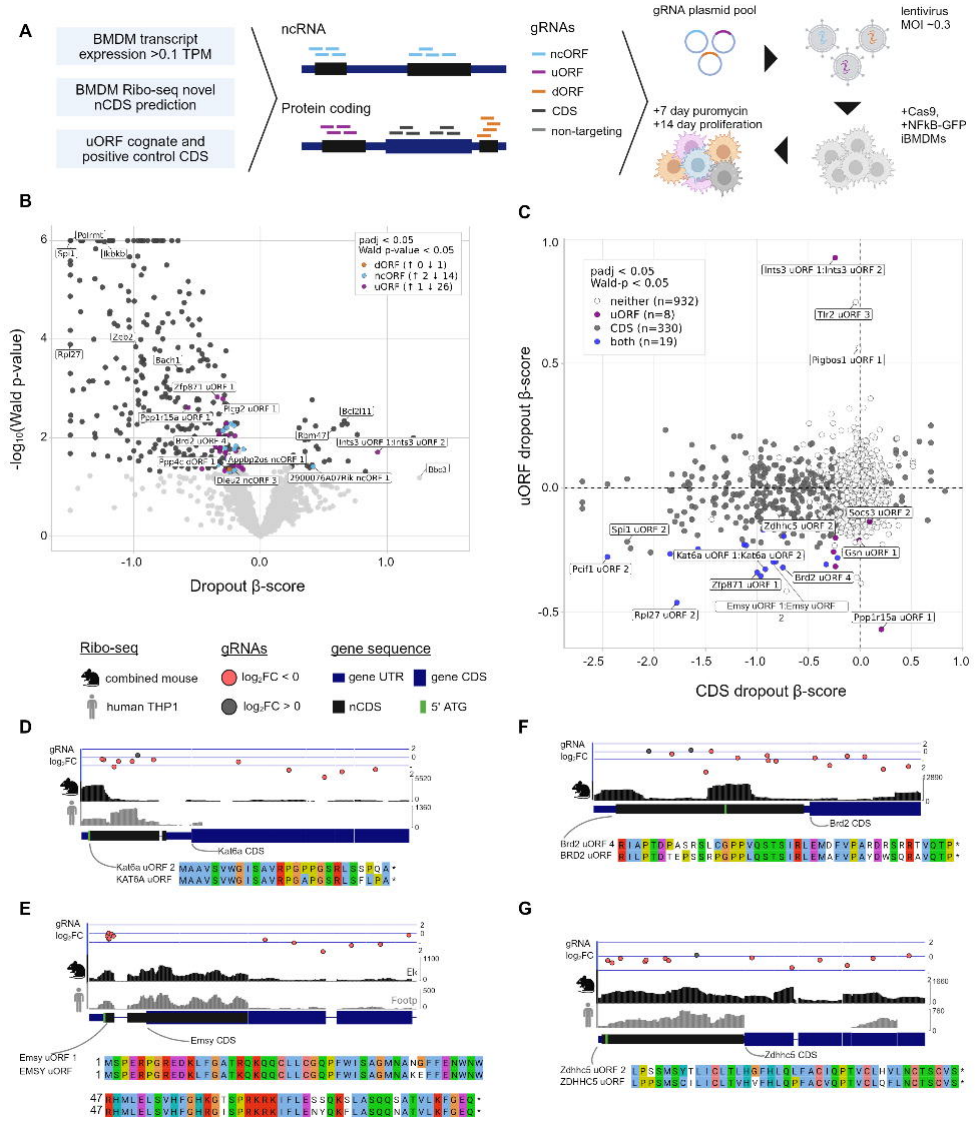
Dropout screen against nCDSs with predicted translation in bone marrow derived macrophages (BMDMs). (A) CRISPRware designed a gRNA library against nCDSs on transcripts expressed in BMDMs (>0.1 TPM). Each ORF was targeted with at least 4 guide RNAs. Where uORFs were targeted, the canonical coding sequence (CDS) was also included. After pooled transduction into immortalized CRISPR-Cas9 BMDMs, cells were selected with puromycin for seven days, and genomic DNA was taken 14 days subsequent to selection. (B) MAGeCK-MLE dropout results for nCDS and canonical CDS hits from the pooled CRISPR dropout screen. ORFs with padj < 0.05 and Wald-p-value < 0.05 were considered significant hits. To enhance visualization, beta scores < −1.5 were capped at −1.5 and Wald p-values < 1E-6 were set to 1E-6. Full values are available in the associated table. (C) Comparison of CDS and uORF effects identifies genes where disrupting the upstream ORF and/or the canonical CDS contributes to fitness. (D-G) NCDS loci illustrating gRNA dropout (mean from triplicates), Ribo-seq support, ORF structure, and conservation with human sequence. Mouse Ribo-seq is combined from all studies (Figure 1). Human THP-1 Ribo-seq was mapped to Hg38 and lifted to mouse Mm39. (D,E,F) In each case the uORF and cognate CDS were significant dropout hits (F) Brd2 did not have signal from lifted THP1 tracks. (G) For Zdhhc5 the uORF passed our significance threshold (padj = 2.3E-10, Wald p-value = 9E-3) but the cognate CDS did not (Wald p-value > 0.05). TPM: Transcript per million; iBMDM: immortalized bone marrow derived macrophages; MOI: multiplicity of infection

Among the most strongly supported examples were interferon-induced loci annotated as pseudogenes, including ENSMUSG000000121501 (Gvin-ps7) and Gm12250. Despite their pseudogene annotation, these loci exhibit uniquely mapped bulk RNA and robust Ribo-seq signals across their dominant ORFs, alongside proteomic evidence of translation in macrophages (Fig. 2G, Supp. Fig. 3A,B). We also identified a cluster on chromosome 13 in which multiple transcripts annotated as long noncoding RNAs neighbor the protein-coding zinc finger gene Zfp825. Once again, these loci showed convergent evidence of translation: uniquely mapped Nanopore reads, bulk RNA and Ribo-seq coverage, unique PSMs, and predicted KRAB/zinc finger domains consistent with ZFP-like proteins (Supp. Fig. 4A-C). Even after strict FDR filtering, PSMs can be spurious so we implemented a set of stringent post-FDR filters based on recent guidance and included plots of “excellent” graded PSMs (Methods) to support these translation events^14^ (Fig. 2H, Supp. Fig. 4C). Together, these cases argue that a subset of “noncoding” or “pseudogene” annotations reflect genuine protein synthesis, supported by independent readouts spanning transcription, ribosome occupancy, and peptide detection.

**Figure 4.**
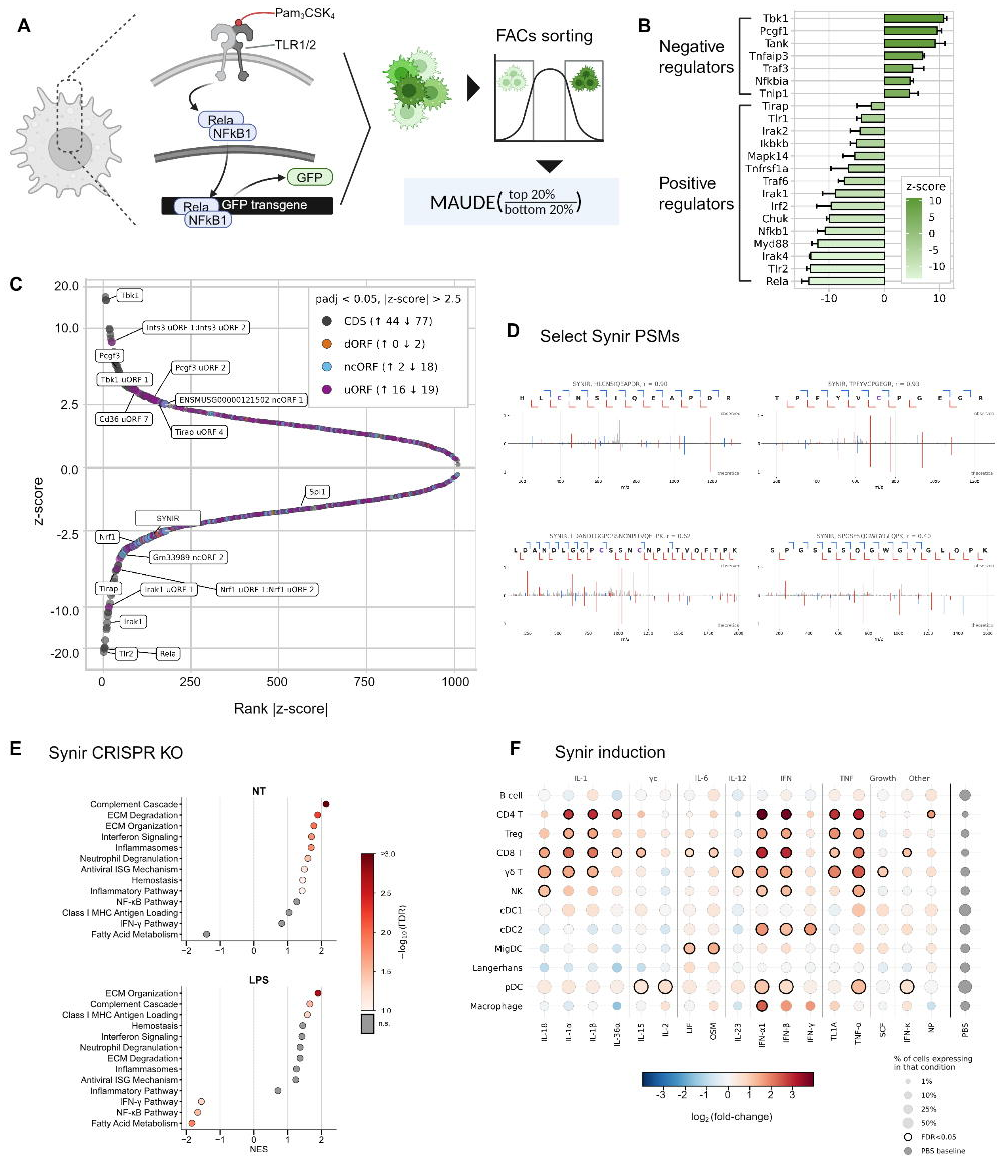
**TLR1/2-NF-κB sorting screen results**. (A) Activation of the Tlr1/Tlr2-NFkB inflammatory cascade induces GFP production in NFkB-GFP immortalized bone marrow derived macrophage (iBMDM) line. Cells were sorted into GFP high (top 20%) and GFP low (bottom 20%) buckets by GFP expression after stimulation with Pam_3_CSK_4_. Guide RNA and gene level z-score were calculated with MAUDE (Mean Alterations Using Discrete Expression). Negative z-scores indicate positive regulators of NFkB signaling, positive z-scores indicate negative regulators. (B) The MAUDE calculated z-scores of known positive and negative regulators of Tlr1/Tlr2-NFkB signaling cascade. (C) ORFs are considered hits if the direction of the effect is the same for all replicates, the absolute values of the combined z-score are > 2.5 and the padj. < 0.05. nCDS combined z-scores across triplicates calculated with Stouffer’s method. Z-scores are log transformed to improve visualization. (D) Peptide spectra that map uniquely to SYNIR, nowhere else in the genome, passed FDR filters, and scored “excellent” according to the heuristic scorer (Methods). (E) Preranked GSEA of genes ranked by DESeq2 Wald statistic (SYNIR KO vs. negative controls) in untreated (NT, top) and LPS-stimulated (bottom) BMDMs. Shown are gene sets with FDR < 0.10 in at least one condition. The x-axis gives the normalized enrichment score (NES); positive values indicate enrichment among genes upregulated in KO. Dot colour gives −log10(FDR), capped at 3.0. Grey dots are not significant in that condition (FDR ≥ 0.10). FDR q-values were computed from 1,000 gene-label permutations. (F) Dot plot of Synir (ENSMUSG00000120992 or D17H6S56E-5) in the mouse Immune Dictionary (lymph node after in vivo cytokine injection). Rows are the 12 cell types of the original annotation; columns are the 17 cytokines that change D17H6S56E-5 significantly in at least one cell type, grouped by ligand family, followed by a PBS column showing baseline detection. Dot colour is the logFC versus PBS from edgeR quasi-likelihood F-tests on per-mouse pseudobulks (design ∼ batch + cytokine). Black rings mark FDR < 0.05, with Benjamini-Hochberg correction within each cell type across all cytokine tests of the eight tested mouse ERV-derived genes.

While proteomic support provides especially compelling evidence for translation, we emphasize that shotgun mass spectrometry remains biased toward abundant, stable proteins and is expected to miss many translated products - particularly small, lowly translated, and unstable proteins. In line with this limitation, while our pipeline reidentified multiple previously published translated ORFs with Ribo-seq (Fig. 1G, Table S4), we did not identify confident PSMs that matched the predicted proteins from any of these 10 nCDSs. Thus, proteogenomic integration serves primarily to prioritize a subset of translation events, rather than to define the full scope of biologically meaningful noncanonical translation.

### A pooled macrophage essentiality screen identifies ORFs that contribute to cellular fitness

To functionally prioritize nCDSs, we designed a targeted CRISPR library against candidates with evidence of transcription and Ribo-seq translation in macrophages (Table S23) using the CRISPRware guide RNA library design package^76^. We targeted 2,165 uORFs and 63 dORFs across 984 protein-coding genes and 1,093 ncORFs across 400 ncRNAs and pseudogenes. In the analysis we consider each ORF that was targeted by at least 4 independent guide RNAs (gRNAs). In some cases, ORFs overlapped due to out-of-frame translation or alternative splicing and gRNAs could not be uniquely associated with a single target; in these cases the co-targeted ORF IDs were concatenated, e.g. Emsy uORF1:Emsy uORF2, and treated as a single targeted unit. For genes harboring targeted uORFs, we additionally included gRNAs against the cognate downstream CDS to enable paired comparisons at shared loci. GRNAs were cloned into lentivirus and transduced at low multiplicity of infection into our previously published immortalized bone marrow-derived macrophage lines, which constitutively express Cas9 and carry an NFκB-GFP reporter cassette^77^ (Fig. 3A, 4A). Following selection and outgrowth in three distinct clonal lines, we quantified changes in guide abundance and inferred gene-level effects on cellular fitness using MAGeCK-MLE^78^.

Encouragingly, an inspection of the top gene hits included common essential genes such as *Polrmt* and Rpl27(depmap.org)^79^ as well as genes that have demonstrated essentiality specifically in macrophages: *Spi1*^80^, *Bach1*^81^, *Ikbkb*^82^, and *Zeb2*^83^. On the other hand, pro-apoptotic *Bcl2l11*^84^ and tumor suppressor *Rbm47*^85^ genes were significantly enriched targets (Fig. 3B). To minimize false positives, we defined confident hits using dual criteria (permutation padj < 0.05 and Wald p-value < 0.05), a threshold that proved sufficiently stringent to exclude genes with described effects on viability, such as the pro-apoptotic factor *Bbc3*, from the high-confidence set (Fig. 3B; Tables S24,S25). Two previously reported nCDSs, Dleu2 ncORF3 and Ppp1r15a uORF1 (Fig. 1G, Fig. 3B, Table S4), were recovered as significant dropout hits, nominating a potential role in macrophage viability in addition to previously ascribed functions^12,34^.

Next we leveraged the paired design to compare uORF-targeting gRNAs with gRNAs against cognate CDS regions: in 19 loci, disruption of either the uORF or the CDS significantly impaired viability, whereas in 8 loci, uORF targeting produced a significant dropout phenotype in the absence of a significant CDS effect (Fig. 3C). We examined four representative uORF hits - Emsy uORF 1, Kat6a uORF 2, Zdhhc5 uORF 2, and Brd2 uORF 4 - and observed consistent negative log fold changes across their targeting guides, consistent with contribution to macrophage fitness (Fig. 3D-G). Each of these uORFs is conserved in the orthologous human gene, suggesting evolutionary constraint on the translated element. In three of the cases, published Ribo-seq reads from the human monocytic THP-1 line, when lifted to mouse genome build mm39 coordinates, recapitulated the ribosome occupancy patterns observed at the corresponding mouse uORFs (Fig. 3D,E,G), indicating conserved translation across species and strengthening the case that these uORFs are important elements in myeloid biology broadly.

### A TLR1/TLR2-NFκB sorting screen reveals noncanonical ORFs that tune innate immune signaling

The same transduced cells were used for an orthogonal screen that directly reads out NFκB pathway activity. At day 14, we stimulated cells with Pam_3_CSK_4_, a proinflammatory bacterial lipopeptide ligand that binds the TLR1/TLR2 heterodimer, triggering canonical NFκB activation and inducing GFP expression via an NFκB-GFP expression element^77,86^. We then performed fluorescence-activated cell sorting to isolate GFP-high and GFP-low populations and sequenced guide-DNA in each bin, enabling identification of nCDS and CDS elements that influence this signaling pathway (Fig. 4A).

We analyzed each replicate independently using MAUDE^87^ to estimate a z-score for each targeted element, then filtered candidate hits to require directional consistency across replicates (all z-scores either positive or negative for a given ORF), after which replicate-level information was aggregated (Methods). We defined significant NFκB modulators as those with |z-score| > 2.5 and padj < 0.05 (Fig. 4C; Tables S26-S28). Many top-scoring hits corresponded to established regulators of TLR-NFκB signaling, supporting assay fidelity (Fig. 4B). Notably, the thresholds remained conservative enough to exclude several likely pathway components that did not meet significance, including Nfkb2 (z-score = 2.21, padj = 0.068), Traf1 (z-score = −2.19, padj = 0.069), and Irak3 (z-score = −2.05, padj = 0.080), indicating that the reported set reflects stringent filtering (Table S28).

To assess whether generalized dropout phenotypes could present as NFκB regulators, we compared effect profiles between the fitness and sorting screens. Overall, the two hit sets were notably independent, indicating that most NFκB modulators were not simply genes required for growth in culture, and conversely that viability hits did not significantly bias GFP output (Supp. Fig. 5A). An exception was a short, 14-codon uORF in Ints3, which emerged as a hit in both assays despite the absence of a corresponding phenotype for the cognate Ints3-CDS, suggesting a locus where a small translated element may exert disproportionate regulatory impact (Supp. Fig. 5B). Conversely, analysis of Irak1 uORF1, our top nCDS hit in the sorting screen, highlighted an important caution for interpreting short-ORF perturbations: the highly active gRNAs substantially overlapped the embedded microRNA mir-718, emphasizing the need to consider local genomic context and to integrate multiple lines of evidence when attributing phenotypes to a given ORF (Supp. Fig. 5C).

**Figure 5.**
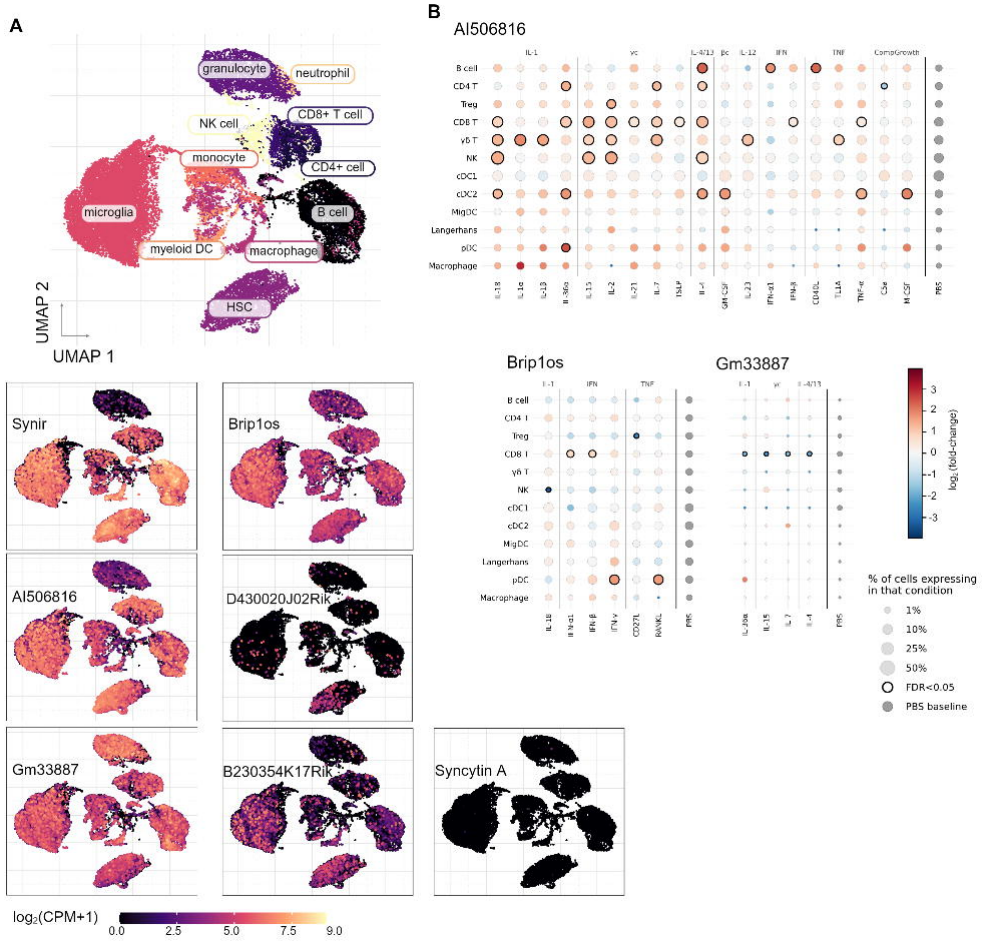
**Env-derived gene expression and induction in immune cells**. (A) Tabula muris expression by immune cell type. Synir (596 codons) (ENSMUSG00000120992/D17H6S56E-5), AI506816 (645 codons), and Gm33887 (641 codons) are predicted to be translated to produce full transmembrane proteins with classic retroviral envelope signatures. Brip1os (282 codon ORF) and D430020J02Rik (226 codon ORF) encode truncated viral proteins with N-terminal signal peptides, but lack the transmembrane viral protein domain. B230354K17Rik harbors a translated 493 codon ORF with homology to Friend virus susceptibility protein 1 - a protein thought to arise from retroviral gag protein endogenization. Syncytin A, a fusogenic protein derived from retroviral envelope endogenization with expression restricted to placental trophoblasts, and is not translated in adult immune cells. (B) AI506816, Brip1os and Gm33887 responses to cytokines across lymph-node cell types in vivo. Dot plots for the lncRNAs AI506816 (top), Brip1os (bottom left) and Gm33887 (bottom right) in the mouse Immune Dictionary (lymph node after in vivo cytokine injection). Rows are the 12 cell types of the original annotation; in each panel, columns are the cytokines that change that gene significantly in at least one cell type, grouped by ligand family, followed by a PBS column showing baseline detection. Dot colour is the logFC versus PBS from edgeR quasi-likelihood F-tests on per-mouse pseudobulks. Black rings mark FDR < 0.05, with Benjamini-Hochberg correction within each cell type across all cytokine tests of the eight tested mouse ERV-derived genes. (C) Western blot (WB) results HEK293s transfected with either SEMR-V5 overexpression vector or control (CTL) vector. WB was performed with v5 antibody on, Left: cell lysate; Middle: V5 immunoprecipitation (IP) of cell lysate; Right: v5-IP of supernatant. Expected size of SEMR-V5 is ∼34 kDa with variable glycosylation expected.

Finally, we highlight two instructive uORF patterns that illustrate how nCDS elements can diverge from their cognate CDSs: a conserved uORF in the transcription factor Nrf1 gene produced a stronger effect on NFκB-GFP output (z = −5.1, padj < 4×10^-6^) than the Nrf1 CDS (z = −3.2, padj < 9×10^-3^), and Tirap uORF4 (z = 2.7) exhibited an effect in the opposite direction of its cognate CDS (z = −5.3) (Supp. Fig. 5 D,E). Together these examples suggest that uORFs can exert measurable impact on a critical signaling pathway, with evidence that they sometimes exhibit effects as profound as, or opposite to, their cognate coding sequence.

### A syncytin-like envelope protein - SYNIR - is implicated in NFκB-regulation

One sorting-screen hit was a 596-codon nCDS on an uncharacterized long noncoding RNA identified as ENSMUSG00000120992 (or D17H6S56E-5) (z-score = −2.8, padj = 0.024). In addition to being a sorting screen hit, translation of this lncRNA was supported by multiple independent lines of evidence: strong Ribo-seq signal across the ORF, robust transcription in macrophages, and 14 unique peptide-spectrum matches that map to the predicted protein sequence (Table S19,S20), including multiple “excellent” grade PSMs that are unique both genomically and transcriptomically (Fig. 4D).

We examined predicted protein domains and discovered protein architecture consistent with a retrovirus-derived envelope protein, including a signal peptide, an extracellular surface unit (SU), and a transmembrane domain (TM) (Supp. Fig. 5F,G). Together, these features suggested that the encoded protein is a full-length, membrane-spanning envelope-like (Env) protein. To place this gene in context, we performed a BLASTp search against mouse and human proteins and identified human Syncytin-1 as the closest characterized homolog (E < 5e^-10^). Syncytins are endogenized retroviral genes that are required for formation of the placenta^40,41,88^. Given the structural similarity to Syncytin-1 and the impact on NFκB signaling, we renamed this protein SYNIR (SYNcytin-like Immune Regulator; gene name: *Synir*), used hereafter. A closer inspection of SYNIR uncovered remarkable conservation of hallmark features with Syncytin-1. These included canonical cysteine motifs consistent with disulfide-stabilized SU/TM organization, a predicted furin cleavage site that would separate SU and TM, and motifs indicating a homotrimeric interface and an immunosuppressive domain (ISD) (Supp. Fig. 5G). These shared structural and functional signatures support interpreting SYNIR as a retrovirus-derived Env protein that has been retained and repurposed in the mouse genome. We next tested whether SYNIR functionally contributes to inflammatory signaling in order to confirm its role as a positive NFκB regulator. We disrupted the SYNIR protein using three independent gRNAs in our Cas9 iBMDM line and performed RNA-seq at baseline and following stimulation with the inflammatory ligand LPS (Tables S29, S30). At baseline, gene pathways related to immune function were enriched, including NFκB and IFNγ (albeit not at FDR <0.1). Following 6 hour LPS treatment, SYNIR disruption again altered the transcriptional state with both NFκB and IFNγ pathways significantly down, in agreement with the screen results. Collectively, these data establish *Synir* as a translated, retrovirus-derived Env protein and implicate SYNIR protein in NFκB-signaling in macrophages.

### Updated scRNA-seq references expand the landscape of envelope-like translated loci across immune and tissue contexts

The identification of SYNIR as a protein of retroviral origin encouraged us to search for additional Env translation events. Across the aggregated Ribo-seq predictions, we identified four lncRNA loci with ORFs whose predicted proteins exhibit Env features: *ENSMUSG00000120992* (*Synir*), *D430020J02Rik*, *Brip1os*, and *AI506816*. One additional Env gene, *Gm33887*, was previously predicted in the RefSeq proteome database (it is annotated as a lncRNA in GENCODE, and our pipeline did not identify it as translated). Because envelope-derived genes are typically discussed in the context of placental formation and fetal development^89^, we sought a broad and unbiased view of expression across tissues and immune lineages. We therefore reanalyzed Tabula Muris^90^, Tabula Senis^91^, and the mouse Immune Cytokine Dictionary^92^, single-cell RNA-seq references using an updated gene annotation that includes the aforementioned Env genes and other missing lncRNAs. This reanalysis provides resources for interpreting lncRNA specificity across adult tissues at three ages (3-,18-, and 24-months; >110,000 total quality filtered cells) and in immune cells treated with dozens of cytokines and can facilitate lncRNA investigations beyond the present study (Data Availability).

In these updated scRNA-seq embeddings, the Env lncRNA loci showed distinct and nonredundant expression patterns across immune subsets and tissue contexts (Fig. 5A; Supp. Fig. 6A-F), in contrast with Syncytin-A, which demonstrated virtually no transcription in these non-placental cells (Fig. 5A). Across genes, we observed marked differences in both the magnitude of expression and the cell types in which expression concentrates, supporting a diversified landscape of envelope-derived loci with lineage- and tissue-biased expression in adult mice (Fig. 5C; Supp. Fig. 6A-F).

**Figure 6.**
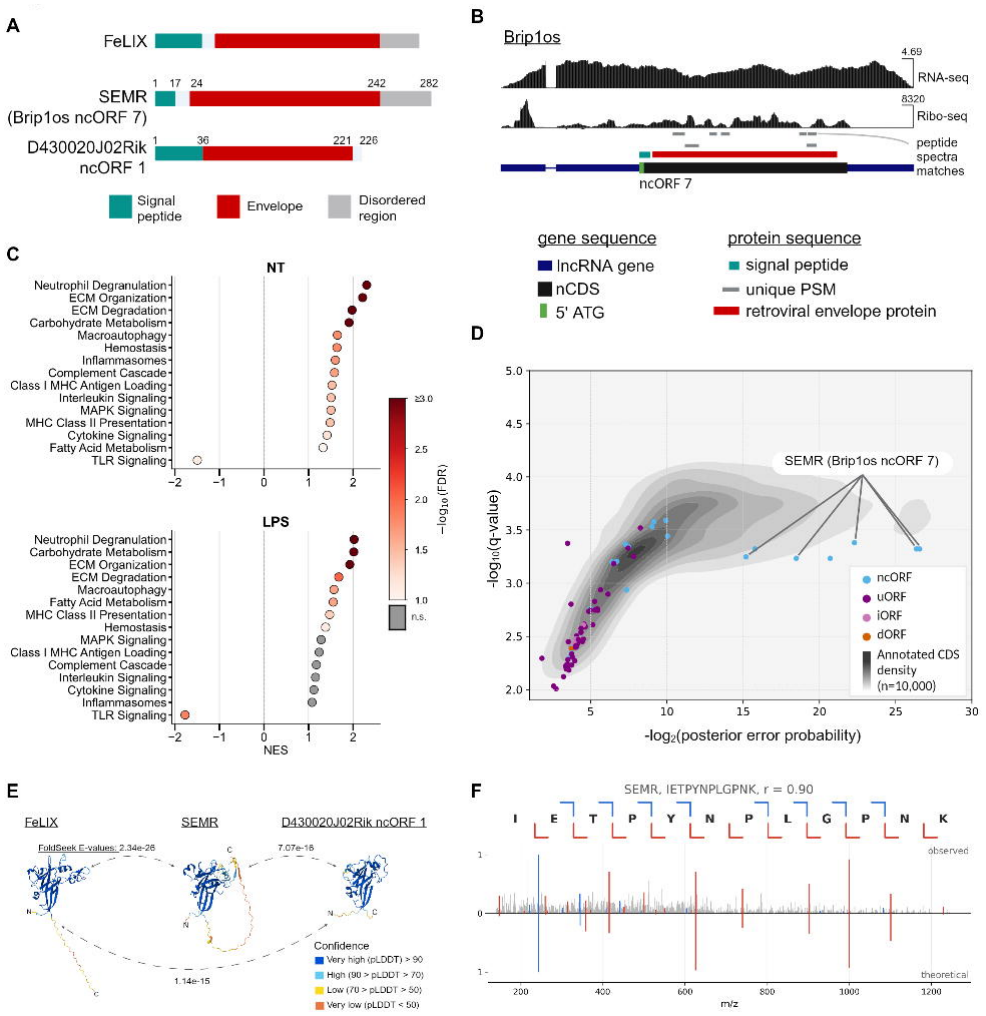
**Brip1os encodes a retrovirus-derived secreted protein, SEMR, in bone marrow derived macrophages (BMDMs) with immunomodulatory function and homology with FeLIX protein**. (A) Predicted domains of Feline Leukemia virus accessory protein (FeLIX), SEMR (Brip1os ncORF7), and D430020J02Rik ncORF 1. (B) Gene set enrichment analysis of Brip1os knockout BMDMs. Preranked GSEA of genes ranked by DESeq2 Wald statistic (Brip1os KO vs. negative controls) in untreated (NT, top) and LPS-stimulated (bottom) BMDMs. Shown are gene sets with FDR < 0.10 in at least one condition. The x-axis gives the normalized enrichment score (NES); positive values indicate enrichment among genes upregulated in KO. Dot colour gives −log10(FDR), capped at 3.0. Grey dots are not significant in that condition (FDR ≥ 0.10). FDR q-values were computed from 1,000 gene-label permutations. (C) Foldseek search reveals close structural homology between each of the three proteins. (D) Unique peptide-spectrum matches (PSMs) from macrophage secretome datasets. PSMs uniquely mapping to the SEMR protein sequence are labelled. (E) Peptide spectra that map uniquely to SEMR, nowhere else in the genome, passed FDR filters, and scored “excellent” (top) or “good” (bottom) according to the heuristic scorer (Methods). NT: no treatment LPS: lipopolysaccharide

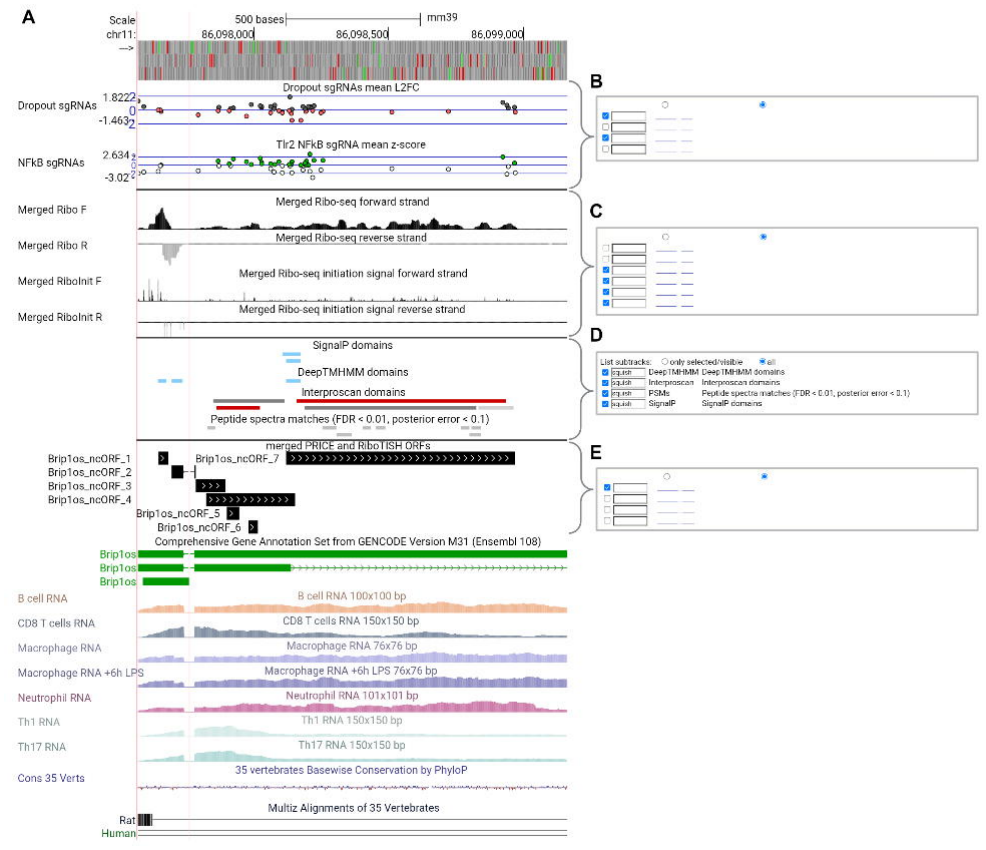

To ask whether these loci respond to immune signals, we tested each gene for cytokine-induced expression changes in the reanalyzed Immune Cytokine Dictionary, in which mice received one of 86 cytokines or PBS^92^. *Synir* and *AI506816* were each detected in roughly 15% of lymph-node cells, *Brip1os* in 7%, and *D430020J02Rik* and *Gm33887* in fewer than 1%. *Synir* was the most cytokine-responsive Env locus (Fig. 4F): it was induced in 51 cell type-cytokine combinations spanning nine of the twelve annotated cell types, with no significant decreases. Type I interferons (IFN-α1 and IFN-β) induced SYNIR in eight cell types, most strongly in CD4+ T cells (log2 fold-changes of 3.8 and 3.7), while IL-1-family (IL-1α, IL-1β, IL-18, IL-36α) and TNF-family (TNF-α, TL1A) cytokines induced it mainly in T and NK cells. *AI506816* instead tracked lymphocyte growth and differentiation signals (Fig. 5B). *Brip1os* and *Gm33887* responded sparsely (6 and 4 significant responses, respectively), and *Gm33887* was repressed rather than induced, by IL-15, IL-7, IL-4 and IL-36α in CD8+ T cells. *D430020J02Rik* showed no significant response in vivo, and *Syncytin-A* and *-B* were too rarely detected to test (4 and 9 counts across all samples).

Because Env-derived genes also persist in the human genome, we asked whether human Env-ERV genes share these cytokine responses, using a cytokine dictionary of human peripheral blood mononuclear cells stimulated in vitro with 90 cytokines^93^. The human syncytins were rarely detected in blood cells (Syncytin-1, *ERVW-1*, 0.03% of cells; Syncytin-2, *ERVFRD-1*, 0.05%), mirroring the near absence of *Syncytin-A* and *-B* in mouse immune cells (Supp. Fig. 7). By contrast, the most broadly expressed human env-class gene, *ERVK13-1* (14% of cells), responded to 71 lineage-cytokine stimuli. Consistent with its interferon inducibility, *Synir*’s cytokine-response profile closely matched human interferon-stimulated genes *Irf7* and *Stat1*. *AI506816* shared IL-4, CD40L and common γ-chain stimuli with the human env-class genes *ERVK13-1* and *ENSG00000287277*. Overall, overlap in the stimuli that induce ERV-derived genes in both species was modest and did not exceed chance once interferon-driven stimuli were excluded, indicating that shared regulation of these loci is largely confined to the interferon response. Together, these data show that, unlike the canonical syncytins, several Env-derived lncRNA loci are actively and selectively regulated in adult immune cells, with *Synir* behaving as an interferon-inducible gene across lymphoid and myeloid lineages and *AI506816* as a lymphocyte cytokine-responsive gene.

### Long-read Nanopore sequencing resolves unannotated retroviral Env gene models and isoforms

Our custom transcriptome incorporated a de novo Nanopore direct RNA assembly from BMDMs^61^, enabling us to identify genes and isoforms that are difficult to resolve with short reads alone. Within this long-read-informed annotation, we identified additional Env ORFs that were absent in GENCODE. Three loci emerged as high-confidence Env genes (Supp. Fig. 8A-D). First, *XLOC000481* corresponds to a novel *Ctse* isoform defined by an alternative second exon that is strongly represented in long-read data relative to canonical *Ctse* isoforms (Supp. Fig. 8A). Short-read junction spanning reads corroborated this isoform structure and relative usage (Supp. Fig. 8B). Second, *XLOC012421* is an unannotated, highly transcribed gene antisense to *Cnbd2* (Supp. Fig. 8C). Third, *XLOC008059* is an extended second exon model of the annotated lncRNA *Gm62403* (Supp. Fig. 8D). Unlike the annotated Env genes above, these three loci exhibit extreme sequence similarity exemplified by an absence of uniquely targeting CRISPR guides. We observed only low Ribo-seq signals at the loci and RiboTISH, but not PRICE, identified the Env ORFs at these loci as translation events. We attribute this to the challenges associated with uniquely mapping ∼30-nt, single-end Ribo-seq reads to repetitive genomic elements. This limitation is overcome with long reads, but we are only able to confidently report on transcriptional, not translational, activity at these loci - a reminder of the limits of current approaches.

### Brip1os encodes a secreted envelope-like protein with FeLIX homology

Among the Env candidates, *D430020J02Rik* and *Brip1os* were notable outliers in predicted protein length: both encoded proteins less than 300 amino acids long, less than half the length of classical Env proteins. Domain prediction shows that these proteins preserve an N-terminal signal peptide and SU-like envelope domain, but lack a transmembrane segment, consistent with secreted, rather than membrane-bound, proteins (Fig. 6A). Mirroring our approach with SYNIR we used BLASTp to search for homologs, but did not find characterized proteins in mouse or human. Next, we used Alphafold 3^94^ to predict structures and found that both produced high-confidence β-sandwich-like architectures. We used Foldseek web server^95^ to search structural databases and identified an ortholog in FeLIX, a previously described feline retrovirus-derived accessory protein with documented roles in modulating the ability of feline leukemia virus to infect host cells^96,97^ (Fig. 6B).

Brip1os-ncORF 7, in particular, emerged as an attractive candidate for functional follow-up because of its widespread and consistent expression across tissues (Supp. Fig. 6E), and as the most prominent nCDSs hit in macrophage secretome proteomics datasets (Fig. 6D). We asked whether the *Brip1os* encoded protein has a measurable regulatory impact in cultured macrophages. Using three independent gRNAs, we disrupted the Brip1os ncORF 7 in iBMDMs and performed RNA-seq at baseline and after 6 hours of LPS stimulation. KO induced extensive transcriptional remodeling in both conditions (Fig. 6C; Table S31,S32), with hundreds to thousands of differentially expressed genes (padj < 0.01, |log2FC| > 1). The impacted genes spanned core macrophage programs, including metabolic and hypoxia-linked regulators (e.g., Hif1a, Bach1), inflammatory and NFκB-associated transcriptional circuitry (e.g., Nfkb1, Junb, Rel), lineage determinants (e.g., Zeb1, Zeb2), and broad chromatin and transcriptional regulators (e.g., Ctcf, Kmt2a). Pathway-level analysis highlighted enrichment pathways related to metabolism and immune activity (Fig. 6C). Based on this protein’s secretion, derivation from a retroviral envelope protein, and impact on cellular metabolism, we name this protein SEMR (Secreted Envelope-like Metabolic Regulator). Notably, despite pronounced changes in this targeted KO experiment, SEMR was not a significant hit in either screen. We then tested whether SEMR was translated and secreted by overexpressing SEMR-V5 in HEK293 cells and Western blotting both cell lysate and supernatant. We found SEMR-V5 in both compartments. These observations imply that SEMR may function as a paracrine signaler, and identifying its requisite receptor(s) presents an intriguing future opportunity.

### An interactive genome browser resource enables rapid community interrogation of noncanonical ORFs and functional evidence

A core theme throughout this study is that confident interpretation of noncanonical translation requires synthesis of multiple independent signals: Ribo-seq evidence, peptide-spectrum support, transcription, protein domain predictions, conservation, and evidence from CRISPR screens. To make these layers directly interrogable by the community, we created an interactive public UCSC Genome Browser^72,73^ session that aggregates the processed data into a navigable format (Supp. Fig. 9A-E).

The session is built on the current mouse reference genome (mm39) and is designed so that any locus can be queried via standard gene-name or chromosome coordinate search. The browser view integrates, in a single view-panel, gRNA effect sizes from both essentiality and sorting screens (with options to view replicate-level or mean-level tracks), aggregate mouse leukocyte Ribo-seq tracks, including separate entries for elongation (cycloheximide treated) and initiation (harringtonine and lactimidomycin). Human THP-1 Ribo-seq tracks mapped to hg38 and lifted to mm39, proteomics evidence including peptide-spectrum matches, and protein feature predictions such as secretion signals and transmembrane domains from Signal6P^98^, DeepTMHMM^99^, protein family domains from Interproscan/Pfam^100,101^ and intrinsically disordered regions from MobiDB-lite3^102^ (Supp. Fig. 9A-D). By lowering the barrier to nCDS evaluation, this resource is intended to accelerate hypothesis generation and prioritization for follow-up studies on the many additional translated ORFs uncovered in this work but not yet functionally characterized. Hosting on the highly reliable UCSC Genome Browser ensures that this work will remain a stable and accessible long-term resource.

## Discussion

Here we present a unified Ribo-seq meta-analysis across mouse leukocyte lineages and implement dual orthogonal CRISPR screens - one measuring cellular fitness, the other NFκB activity - to identify noncanonical coding sequences (nCDSs) with measurable biological activity in macrophages. This strategy nominates uORFs and ncORFs that modulate viability and innate immune signaling, and unexpectedly uncovers multiple retrovirus-derived envelope proteins expressed and translated outside the placenta, including SYNIR, a membrane protein that tunes NFκB-responsive transcription and SEMR, a secreted protein with broad paracrine impact on macrophage gene programs.

### Expanding the landscape of noncanonical translation

Our Ribo-seq meta-analysis, cross-species, and proteogenomic integration substantially expand the catalog of translated noncanonical nCDSs. The pipeline reidentified previously validated nCDSs, providing confidence in the approach. Among the most compelling new cases are loci where convergent multiomics evidence rebuffs existing annotations. Interferon-induced pseudogenes such as Gvin-ps7 and Gm12250 exhibit robust Ribo-seq signal and proteomic support in macrophages, and a cluster of chromosome 13 lncRNAs neighboring Zfp825 show translation of ORFs with predicted KRAB and zinc finger domains. These examples illustrate that relatively large protein coding sequences can be missed even in thoroughly annotated genomes.

Moving beyond cataloging, our dual CRISPR screens reveal that a subset of these translated elements carry measurable biological activity in macrophages. The observation that uORFs can exert phenotypic effects on NFκB signaling that are on par with, or opposite to, their cognate coding sequences, has important implications. Classical models of uORF function emphasize translational repression of the downstream CDS, wherein uORF translation attenuates main-ORF output by sequestering ribosomes and/or triggering nonsense-mediated decay. In most cases, we observed the opposite with CDS and uORF KO, resulting in directionally consistent phenotypic impact (Fig. 3C-G). In these cases, it is possible that gRNAs targeting the 5’ UTR reduce gene transcription, thereby reproducing the phenotype of disrupting the CDS directly. However, the added evidence of cross-species conservation of the uORFs argues that these are bona fide coding sequences producing peptides with independent phenotypic effects. Whether the peptides encoded by these uORFs act in *cis* (e.g., by modulating co-translational folding or localization of the nascent main-ORF protein) or in *trans* (e.g., by interacting with cellular machinery independent of the main ORF) remains an open question that warrants targeted biochemical follow-up.

### Retrovirus-derived proteins as an underexplored class of effector proteins

The most unexpected finding of this study is the identification of widespread transcription and translation of envelope-derived proteins in adult mouse cells. Endogenous retroviruses (ERVs) comprise roughly 8–10% of mammalian genomes, but only a handful of their coding remnants have been functionally characterized, most notably the envelope-like (Env) syncytins, which are required for placental development^40,41,88^. Our data implicate ERV-Env loci as important genes in adult cells and tissues indicating novel roles.

SYNIR (ENSMUSG00000120992) emerged from the NFκB sorting screen as a translated lncRNA encoding a 596-amino acid protein with all the hallmarks of a full-length envelope glycoprotein (Supp. Fig. 5D). Its closest characterized homolog is human Syncytin-1, a fusogenic protein with expression restricted to placental and fetal tissue. SYNIR, on the other hand, is highly expressed in adult myeloid, lymphoid, mammary gland, and large intestine tissue (Supp. Fig. 6B). CRISPR KO of SYNIR altered NFκB pathway in a manner consistent with the sorting-screen linking SYNIR with positive regulation of NFκB activity. While support for SYNIR as a screen hit is supported by our targeted RNA-seq experiment we are intrigued by the fusogenic potential of this syncytin homolog (Fig. 5D). The hallmark of syncytin cell fusion is the generation of multinucleated cells - an important part of placental maturation. Notably, macrophages are also known to form multinucleated cells with pathological function: osteoclasts, Langhans giant cells, and foreign body giant cells^103^. The idea that SYNIR (or other Env proteins reported herein) contributes to this phenomenon is exciting, but support for this possibility is tempered by expression of this gene in HSCs, B cells, and other cell types without documented multinuclear cell formation. Still, we see this as an exciting opportunity for future study.

Notably, the most up- and down-regulated genes in SYNIR knockout: *Ccr5* and *Sut1a1*^104^, respectively, are orthologs to proteins linked to human immunodeficiency virus (HIV) infection in humans. We also notice that SYNIR is highly expressed in mammary gland tissue, a compartment which can facilitate retroviral infection through lactation. Additionally expression is high in myeloid, marrow, and large intestines, all compartments in which retroviral infection might occur. These observations resonate with a broader theme in ERV biology: that endogenized viral genes can be co-opted to modulate host susceptibility to exogenous retroviruses. Other germane examples of this phenomenon include *Fv1*, a gag-derived gene that encodes Friend virus susceptibility protein 1, and which is closely related to an nCDS we identified on the lncRNA *B230354K17Rik*; and Feline leukemia virus susceptibility protein (FeLIX), a secreted Env fragment, known to modulate leukemia virus infection in felines, and which is a structural ortholog of another of our documented Env proteins, SEMR.

Because FeLIX has only been studied in the context of modulating feline leukemia virus infection, we were surprised to find that knockout of SEMR induced striking transcriptional changes affecting hundreds of genes - including master regulators of inflammation (Nfkb1, Rel, Junb), hypoxia (Hif1a, Bach1), and lineage identity (Zeb1, Zeb2). This strong KO response leads us to hypothesize that SEMR is not narrowly involved in modulating viral infection as might be expected from work on FeLIX, and leads us to believe that some ERV proteins can act beyond retroviral infection modulation.

Based on the evidence compiled in this report we suspect SEMR is involved in paracrine signaling: it is secreted as indicated by its signal peptide domain, homology with known secreted protein FeLIX, and is most prominent nCDS-protein in macrophage secretome proteomics. It did not appear as a screen hit, but this is also consistent with paracrine function because in a pooled screen, secreted factors are diluted across the culture, masking signaling effects. Moreover, our updated scRNA-seq analysis shows near-ubiquitous expression of the SEMR host gene Brip1os, across cell types, indicating that this protein may be involved in widespread signaling, beyond myeloid cells. Identification of the cognate receptor(s) for SEMR is a priority for future work and may reveal new axes of immune regulation and beyond.

## Limitations and future directions

Several limitations of this study warrant consideration. First coverage across immune lineages is uneven: macrophages and CD4 T cells dominate the catalog, with fewer datasets representing other cell types. Second, our proteogenomic integration relies on shotgun mass spectrometry, which is biased toward abundant, stable proteins. Many translated products - particularly small, lowly expressed, or rapidly degraded peptides - likely evade detection. The absence of PSMs for 10 previously validated nCDSs in our reanalysis underscores this limitation (Fig. 1G). Third, while we were able to directly test a significant fraction of nCDSs in targeted macrophage screens, the majority of the nCDSs remain uninvestigated. Fourth, distinguishing peptide-mediated effects from translational-regulatory (e.g. uORF regulation of cognate CDS), RNA-mediated effects, or DNA (e.g. enhancer/promoter effects) will require targeted biochemical and genetic dissection. Finally, further characterization of SYNIR, specifically its fusogenic potential, and SEMR, including its impact on other cell types and its associated receptor, provide additional opportunities for further research.

## Resource and data availability

A central goal of this study was to lower barriers to community engagement with noncanonical translation data. The interactive UCSC Genome Browser session we provide integrates gRNA effect sizes, Ribo-seq tracks, proteomics evidence, and protein feature predictions in a single navigable interface. By making these data layers directly accessible, we hope to accelerate hypothesis generation and follow-up studies on the thousands of translated ORFs identified here but not yet functionally characterized. https://genome.ucsc.edu/s/emalekos/Leukocyte_ORF_paper

The updated single-cell RNA-seq annotations, incorporating lncRNAs absent from original Tabula Muris and Tabula Senis releases, further extend the utility of this work for the broader research community.

https://github.com/ericmalekos/scRNAseq

Code associated with this manuscript is available at

https://github.com/ericmalekos/leukocyte_nCDS

https://github.com/ericmalekos/psm-analysis

Sequencing data related to the CRISPR screen: GSE319779

Sequencing data related to RNA-seq: GSE319783

## Acknowledgments

Guide RNA vector cloning, lentivirus production, pooled infection, and subsequent NGS library preparation, as well as targeted CRISPR knockout experiments in SYNIR and SEMR, were performed by UCSC CRISPR Core Director Dr. Alice Devigne. Sorting by GFP intensity was performed by the UCSC Institute for the Biology of Stem Cells Flow Cytometry Core Manager, Dr. Bari Nazario. The authors would like to thank Prof. Angela Brooks for advice and guidance on the project.

S.C. is supported by R35GM137801 from NIGMS, and E.M. is supported by F31AI179201 from NIAID.

## Authorship Contributions

S.C. and E.M. conceptualized and administered the project. E.M. curated and analyzed data, developed software, and managed digital resources. E.M. conducted screen outgrowth experiments. V.S. cultured cells and processed RNA-seq samples. L.S. performed western blot experiments. E.M. drafted the manuscript. S.C. secured funding, supervised the project, and reviewed and edited the manuscript.

## Declaration of generative AI and AI-assisted technologies

Claude Opus 4.6 web interface was used in editing and proofreading of this manuscript. Claude Code Opus 4.8 was used to generate the code in the psm-analysis repository.

## Methods

### KEY RESOURCES TABLE

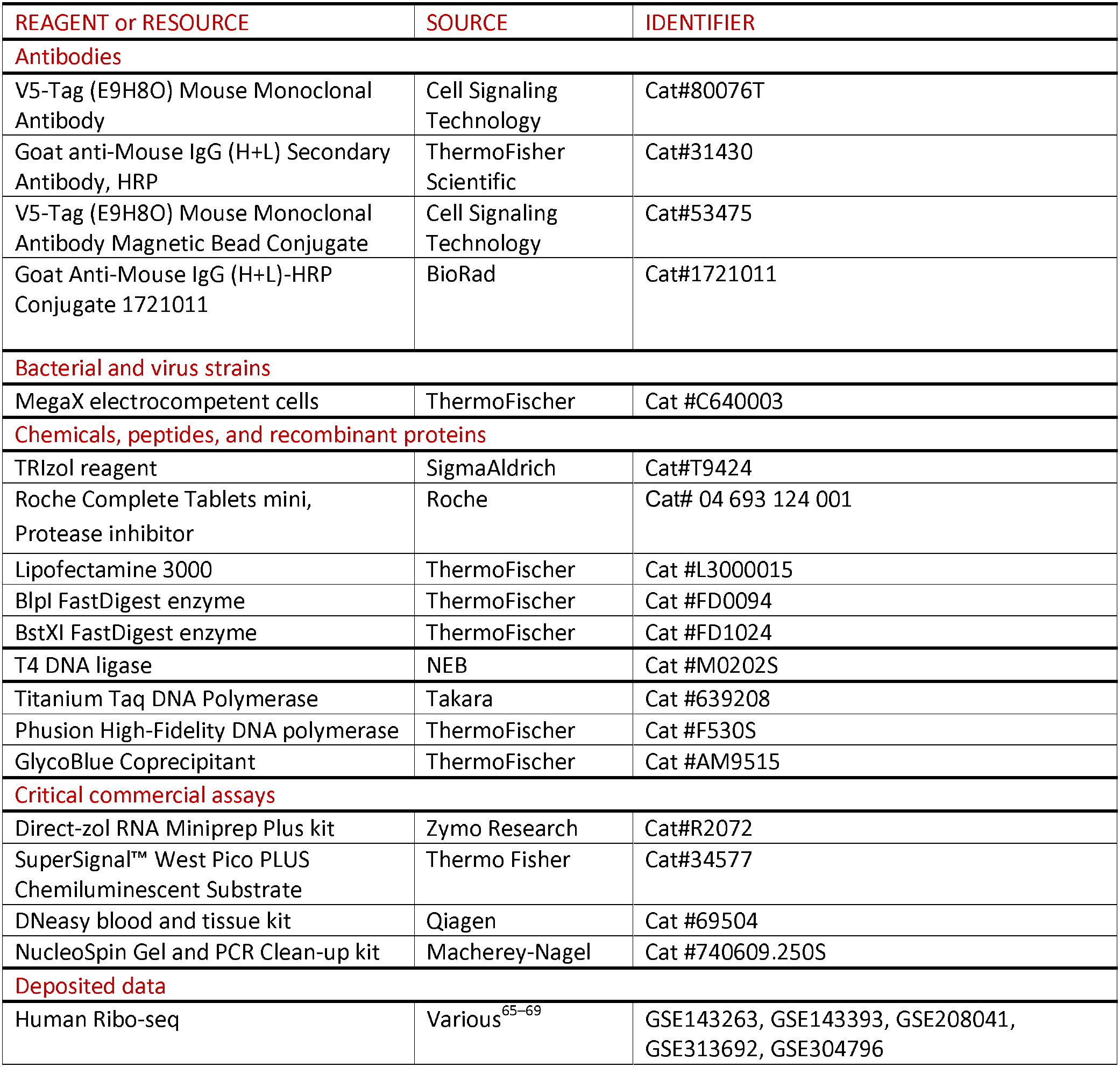
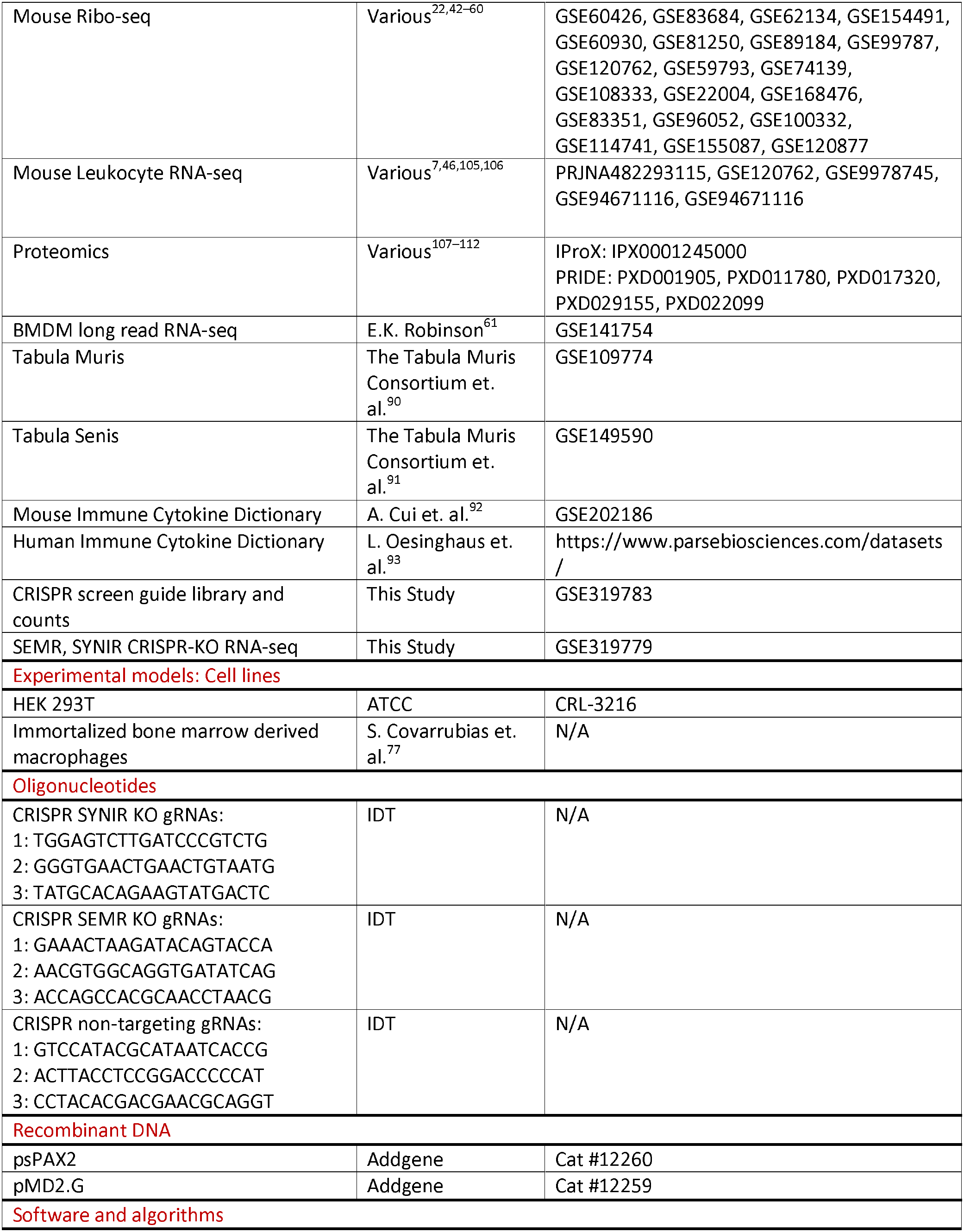
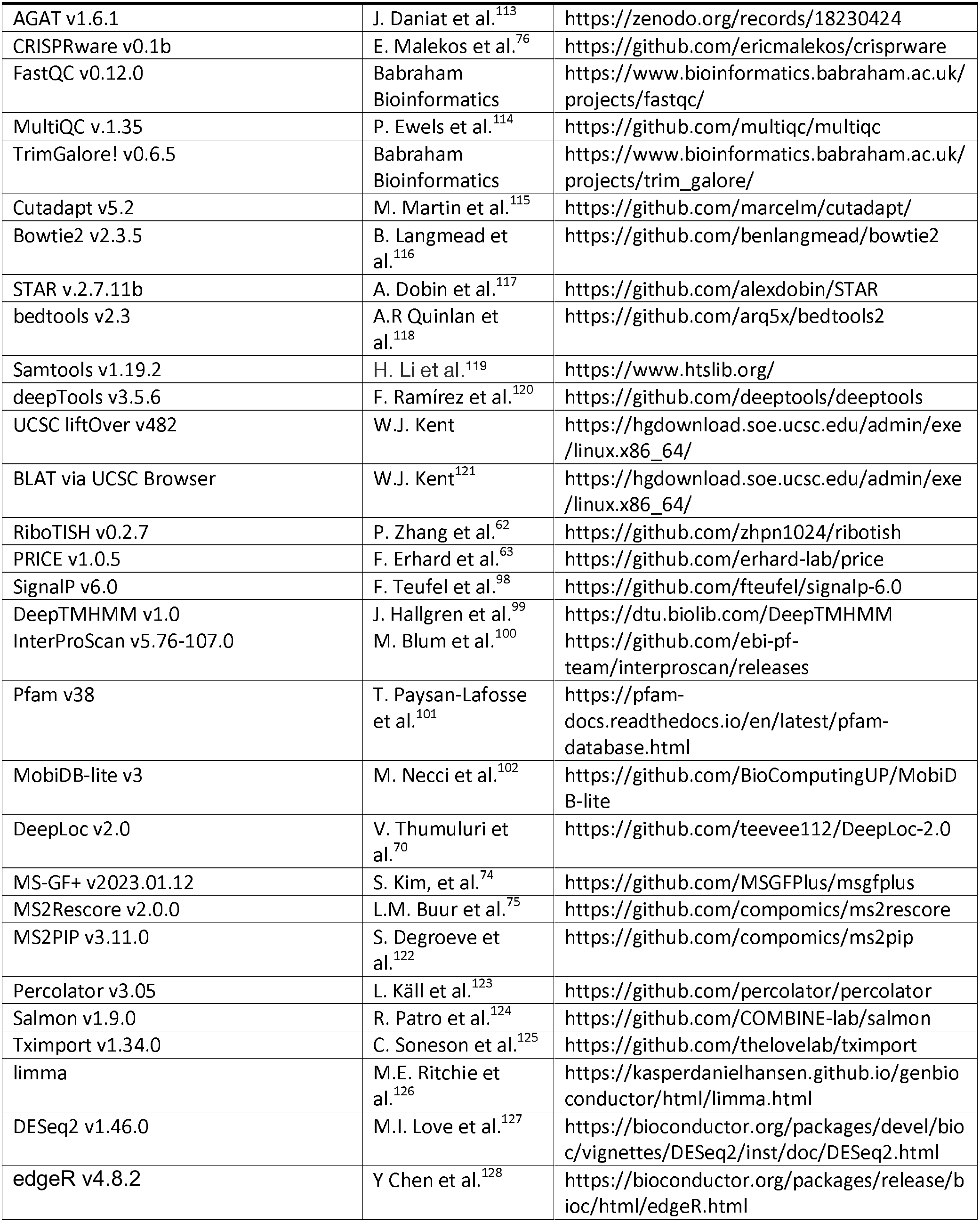
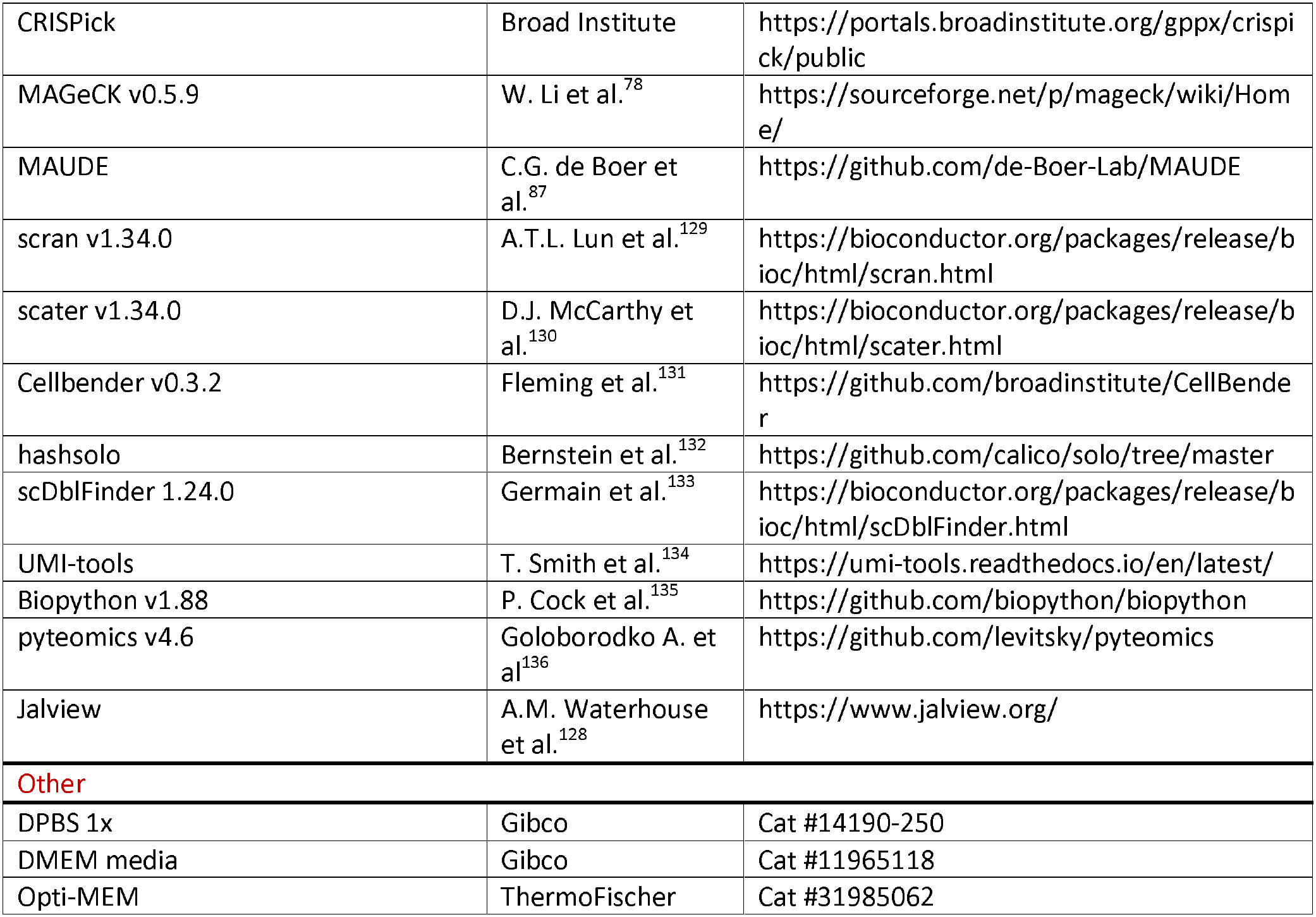

### EXPERIMENTAL MODEL AND STUDY PARTICIPANT DETAILS

#### Immortalized bone marrow-derived macrophages

Clonal lines from our previously published NFκB-GFP+, Cas9+ immortalized bone marrow-derived macrophages (iBMDMs)^77^ were used as replicates in the CRISPR screens. Cells were cultured in DMEM with 10% FBS and 1% Penicillin/Streptamycin at 37 C and 5% CO_2_. Targeted follow up experiments were performed in the same conditions on a single clonal line.

#### HEK293T cells

HEK293T cells were cultured in DMEM with 10% FBS and 1% penicillin/streptomycin at 37 C and 5% CO_2_.

### METHOD DETAILS

#### Ribo-seq data processing

##### Custom gene annotation

Comprehensive gene annotation v31 was downloaded from GENCODE^137^. NCBI RefSeq gene annotation ncbiRefSeq.2020-10-27^138^ was downloaded, and transcripts with transcript_biotype “lnc_RNA” were extracted. We previously generated a long-read de novo BMDM transcriptome assembly^61^. These three files: GENCODE vM31, NCBI Refseq lncRNAs, and BMDM long-read genes were merged with Another GFF Analysis Toolkit’s agat_sp_merge_annotations utility.

##### Mouse Ribo-seq data

GEO datasets were retrieved and FASTQ files were assessed with FastQC and MultiQC. Adapters were removed with TrimGalore!. Mouse tRNA and rRNA sequences were downloaded from GtRNAdb^139^ and RNACentral^140^, respectively, and a bowtie2 index was generated. Trimmed reads were mapped to the t/rRNA index with ‘--very-fast-local’ flag, and unmapped reads were retained. Retained reads were mapped to mouse genome assembly mm39 with STAR with splice sites defined in the custom transcriptome and the following arguments: -- outFilterMultimapNmax 1, --outSAMattributes All, --outFilterMismatchNmax 3, --alignEndsType EndToEnd, --outFilterIntronMotifs RemoveNoncanonicalUnannotated --twopassMode None. Alignments to chrM and nonstandard chromosomes were discarded, and alignments that overlapped GENCODE vM31 snoRNAs were filtered out with bedtools intersect -v -s -f 0.33.

Aggregate signal tracks were formed by separately merging elongating alignments (cycloheximide-treated) and initiating alignments (lactimidomycin/harringtonine-treated) across datasets with samtools merge. Alignments were converted to Bigwig with the deepTools bamCoverage and split by strand. The Ribo-seq signal was not normalized.

##### Human Ribo-seq data

Five human Ribo-seq datasets were analyzed as above, with the following differences. Human tRNA and rRNA sequences were downloaded from GtRNAdb and NCBI RefSeq, respectively. Retained reads were mapped to the hg38 human genome assembly with STAR, with splice sites defined in a merged annotation of GENCODE v49, NCBI RefSeq, and a previously generated a long-read de novo macrophage transcriptome assembly^61^, and the following arguments: --outFilterMultimapNmax 1, --outSAMattributes All, --outFilterMismatchNmax 1, -- alignEndsType EndToEnd, --outFilterIntronMotifs RemoveNoncanonicalUnannotated, -- twopassMode None, --outSAMmultNmax 1. Alignments that overlapped GENCODE v49 snoRNAs were filtered out with bedtools intersect - -s -f 0.33.

For libraries with UMIs, UMIs were extracted to read names with UMI-tools(7 nt for GSE313692, 12 nt for GSE304796, and 6 nt for GSE143393), and 3′ adapters were removed with cutadapt (- e 0.1, -O 3, --discard-untrimmed). A single non-templated base was then clipped from the 3′ end of GSE313692 reads and the 5′ end of GSE143393 reads. Reads were filtered with -q 10, --trim- n, --max-n 0 and a length window of 20-40 nt (20-50 nt for the P1 nuclease-digested GSE313692 libraries). UMI libraries were deduplicated with umi_tools dedup.

THP-1 alignments were lifted to mm39 coordinates with the liftover utility and the hg38ToMm39 chain file provided by the UCSC Genome Browser.

##### ORF calling and filtering

###### ORF calling

Biological replicates within GEO datasets were merged and filtered for alignments in the 27-32 nucleotide length range. RiboTISH’s QC module was run on each library and manually examined for 3-nucleotide periodicity for each set of read lengths. CDSs were called using a hierarchical strategy with both RiboTISH and PRICE at three levels: (1) replicate-level merging within individual experiments, (2) experiment-level merging within each dataset, and (3) global merging across all datasets.

RiboTISH was run with --alt --longest to identify the longest statistically significant CDS with any near-cognate start codon. PRICE was run with default settings, and Benjamini-Hochberg correction was applied to generate adjusted p-values (padj). NCDSs from both tools were retained if they reached padj < 0.01 in at least one hierarchical test and encoded peptides >10 amino acids. ORFs sharing start or stop codons with GENCODE protein-coding genes were excluded to focus on truly novel translation events.

###### Removal of annotated coding sequence fragments

Four annotation rules were applied to each caller’s ORFs separately. Each rule was evaluated on the nCDs’s own genomic blocks against GENCODE. An nCDS was removed if:

1. it was an “ncORF” on a protein-coding host gene.
2. it lay on a protein-coding gene and its blocks fully contained at least three CDS exons of one transcript (note that this could remove long iORFs).
3. it was in frame with an annotated CDS at its own locus, defined as a contiguous overlap of at least 30 nt between an ORF block and a same-strand CDS segment in which the CDS codon phase (from the GTF phase field) equaled that of the ORF. or
4. its start or stop codon coincided with an annotated start or stop codon.

The final nCDS set was generated by merging PRICE and RiboTISH calls, consolidating entries that (1) shared a stop codon and (2) where one amino acid sequence was a complete subsequence of the other, retaining the longer variant.

###### Annotating nCDSs

Amino acid sequences from the final set of novel nCDSs were annotated for functional domains with SignalP6 (signal peptides), DeepTMHMM (signal peptides and transmembrane domains), MobiDB-lite3 (intrinsically disordered regions), and InterProScan5/Pfam (protein families) with default settings. Localization predictions were determined using DeepLoc2.

##### Cross-species ORF comparison

###### Sequence homology: reciprocal best BLASTp hits

For each species, a search database combined its nCDS peptides with all GENCODE proteins translations (human release 49, mouse release M31). Terminal stop symbols were removed and sequences shorter than 5 aa were excluded. Decoy databases of identical size and composition were generated by shuffling the residues of each sequence independently with a fixed random seed.

Databases were formatted with makeblastdb and searched with BLASTp using parameters suited to short peptides: -matrix PAM30 -word_size 2 -comp_based_stats 0 -evalue 10 - max_target_seqs 10, with default PAM30 gap costs (existence 9, extension 1). Human nCDSs were searched against the mouse target and decoy databases, and mouse nCDSs against the human target and decoy databases. For each query, the best hit was the target with the lowest E-value.

A human nCDS *h* and a mouse nCDS *m* formed a reciprocal best hit (RBH) pair when four conditions held: the best hit of h was m, the best hit of m was h, the E-value of h against m was ≤ 10⁻⁵ and the E-value of m against h was ≤ 10⁻⁵. A query whose best hit was an annotated protein could not form a pair.

###### Decoy calibration

For each search direction and E-value threshold t, the false discovery rate of nCDS best hits was estimated as:

● numerator: the number of queries whose best decoy hit was a shuffled nCDS with E ≤ t;
● denominator: the number of queries whose best real hit was an nCDS with E ≤ t.

At t = 10□□ this gave 12 decoy versus 1,614 real best hits for human queries (0.7%) and 15 versus 1,491 for mouse queries (1.0%).

###### Positional conservation by reciprocal stop-codon liftOver

Positional conservation was assessed for all 18,701 human nCDSs and all 22,572 mouse nCDSs. For each nCDS, the stop codon was defined as the last three nucleotides of the ORF in transcript orientation and written as a strand-specific BED interval. Human stop codons were projected from hg38 to mm39, and mouse stop codons from mm39 to hg38, with UCSC liftOver (v482)□□,□□ and the UCSC hg38ToMm39 and mm39ToHg38 chain files. Default parameters were used (minimum match 0.95, so all three codon positions had to map). Stop codons of 14,131 human (75.6%) and 19,439 mouse (86.1%) nCDSs were projected. We anchored on the stop codon because start codon assignment varies between callers and datasets owing to near-cognate initiation, whereas a shared stop codon fixes the reading frame.

Each projected stop codon was compared with the stop codons of nCDSs from the other species on the same chromosome and strand, using the 5′ base of each codon in transcript orientation. A human-mouse pair was called positionally conserved when the projected stop codon coincided exactly with the partner’s stop codon in both directions.

###### Integration and pair annotation

RBH and positional pairs were combined into 2,485 human-mouse pairs (Fig. 2B; Table S[X]). Of these, 857 were supported by both criteria, 1,158 by position only and 470 by sequence only.

To assess whether pairs preserved a single reading frame, human and mouse ORF nucleotide sequences (start to stop codon) were aligned globally with the Biopython PairwiseAligner (match +2, mismatch −3, gap opening −5, gap extension −2). A pair was classified as single reading frame when every internal gap had a length divisible by 3.

###### ORF conservation

For uORF alignment panels (Fig. 3D-G, Supp. Fig. 5B-E), mouse nCDS nucleotide sequences were BLATed against the human genome hg38 build on the UCSC Genome Browser. In cases where an nCDS was identified on the orthologous human gene, the translated sequences were aligned and visualized in the Jalview application.

##### Proteomics

###### Proteomics data

**Database construction**

Cell-type-specific proteome databases were generated for macrophages, B cells, and T cells by combining nCDSs with GENCODE vM31 proteins from transcripts expressed (TPM > 0) in the following RNA-seq datasets:

● Macrophages: PRJNA482293, GSE120762, GSE99787
● T cells: GSE94671
● B cells: GSE94671

For immunopeptidomics analysis of the mouse tissue atlas, a pan-tissue database containing all GENCODE proteins plus all nCDSs was used. MaxQuant common contaminant sequences were appended to all databases.

**Peptide assignment**

Mass spectrometry data from IProX^141^ and PRIDE^142^,

● Macrophages: IPX0001245000
● Macrophage secretomes: PXD001905, PXD011780, PXD017320, PXD029155
● B cells: PXD022099
● T cells: PXD022099
● Immunopeptidomics: PXD008733

Were searched against cell-type-matched databases using MS-GF+ with a reverse-decoy database for false discovery estimation. Carbamidomethylation of cysteine was set as a fixed modification and methionine oxidation as a variable modification. For TMT-labeled dataset PXD029155, TMT (+229.16) was included as a static modification on N-termini and lysines.

**Rescoring and filtering**

MS2Rescore was used to rescore MS-GF+ output with ms2pip and percolator. Trained models were selected based on fragmentation method: “HCD2019” for HCD datasets, “Immuno-HCD” for immunopeptidomics HCD, “TMT” for TMT-labeled samples, and “CID” for all other acquisition types. PSMs matching GENCODE proteins were excluded, and retained PSMs matched at least one nCDSs. NCDS PSMs with q-value < 0.01 and posterior error probability (PEP) < 0.1 were reported.

###### Spectral quality grading of PSMs

To evaluate how well each spectrum supports its assigned peptide, independently of the statistical identification (q-value < 0.01 and PEP < 0.1), we graded every PSM supporting a noncanonical ORF of interest . Grades were assigned with an automated implementation of the manual PSM validation procedure of the TransCODE Consortium^14^.

Seven criteria, each corresponding to an element of the manual procedure, were evaluated: (1) at least 90% of cleavage sites covered; (2) no two consecutive cleavage sites uncovered; (3) the b2 ion observed, or a complete y series (all L-1 y ions); (4) precursor mass error ≤ 12 ppm (≤ 13 ppm for PXD029155, which showed a systematic offset of about -12 ppm); (5) no more than two oxidized Met; (6) the base peak explained; and (7) no unexplained peak with ≥ 50% of the base peak intensity. Grades were assigned in order of precedence: caution, if an unexplained peak reached ≥ 60% of the base peak while coverage was below 85% (a dominant unexplained peak on an incomplete fragment ladder, consistent with co-isolation or misassignment); low information, if coverage was below 55%; excellent, if all seven criteria were met; and good otherwise. Excellent, good and low information correspond to the categories of Deutsch et al. Their false positive and close-but-false-positive categories, and criteria that require expert judgement (fragment intensities at Pro, and plausible alternative residues for unannotated peaks within ladder gaps or beyond the last matched ion), were not automated; caution marks spectra that warrant such scrutiny. Following the manual procedure, in which the PSMs of a peptide are examined until one provides excellent evidence, each peptide was assigned the best grade among its PSMs.

For display, fragment intensities were predicted with ms2pip using the model matching each dataset’s fragmentation (HCD2019, CID, Immuno-HCD or TMT). The Pearson correlation between base-peak-normalized observed and predicted intensities over all predicted singly charged b and y ions (observed intensity set to zero for unmatched ions) is reported with each spectrum but was not used for grading. Grading and plotting used custom Python code built on pyteomics available at https://github.com/ericmalekos/psm-analysis.

###### Genome search for loci encoding PSM peptides

Peptides supporting the ncORFs of interest (env-genes, zinc finger proteins, and pseudogenes) were required to map only to nCDSs in the proteomics search database. However, a peptide unique within it could still be encoded elsewhere in the genome. We therefore searched the genome for every locus that could encode each peptide. Each distinct peptide with at least one PSM passing q < 0.01 and PEP < 0.1 (96 peptides from the 11 noncanonical ORF loci of interest) was searched against the six-frame translation of the GRCm39 primary assembly (GENCODE; 22 chromosomes and 39 unlocalized or unplaced scaffolds). Every exact, full-length occurrence of each peptide was recorded. Because Ile and Leu are isobaric and cannot be distinguished by MS, they were treated as identical during matching, and occurrences that differed from the peptide only at Ile/Leu positions were counted as matches and flagged.

Each peptide was classified as unique to its target (on-target loci only), target plus off-target (on-target and at least one off-target locus), off-target only, or no genomic match. Because the genome is searched as an unspliced sequence, a peptide whose coding sequence spans an exon-exon junction does not match its own locus; no genomic match therefore does not count against an identification.

To assess whether matched loci was likely to produce the peptide, each match was compared with the GENCODE M31 comprehensive annotation and classified as exonic-sense (overlapping an exon of an annotated transcript on the same strand as the peptide’s reading frame), exonic-antisense, intronic (within an annotated gene but outside its exons) or intergenic. Annotated genes with an exonic-sense match were reported as the annotated genes consistent with each peptide (Table S).

##### Bulk and long-read RNA sequencing

###### short read RNA-seq

RNA expression for cell-type comparison (Figure 2A) was quantified with Salmon (-- validateMappings --seqBias --gcBias) against GENCODE vM31 using datasets GSE60927, GSE210222, GSE88987, GSE120762, PRJNA482293, GSE141745. Transcript-level quantifications were aggregated to gene-level counts with tximport, log-transformed, and batch-corrected using limma’s removeBatchEffect function.

###### RNA-seq visualization tracks

For locus-level visualization (e.g., Figure 2C,D), paired-end RNA-seq reads from GSE94671 (100 bp), GSE195678 (101 bp), GSE201870 (150 bp), GSE210419 (150 bp), and GSE120762 (76 bp) were aligned with STAR, with multimapping reads discarded (--outFilterMultimapNmax 1 --twopassMode Basic). Alignments were converted to CPM-normalized BigWig tracks with deepTools bamCoverage.

###### Long read RNA-seq

We previously published Nanopore direct RNA data from wild-type C57BL/6 BMDMs (untreated and 6-hour LPS) that were basecalled and aligned to mouse build mm10. We lifted the alignments to mm39 with the liftover utility and mm10.mm39.chain file provided by the UCSC Genome Browser (https://hgdownload.soe.ucsc.edu/downloads.html) for visualization.

##### CRISPR screens

###### gRNA Library design

In order to generate a gRNA library specific to nCDSs relevant to macrophage biology we used the utilities and workflow from the CRISPRware software package. Novel ORFs were filtered to include only those on RNAs expressed at >0.1 TPM in BMDM-specific RNA-seq datasets (GSE60930, GSE81250, GSE89184, GSE99787, GSE120762) and called at the individual condition level (i.e., before merging across conditions) in at least one macrophage Ribo-seq dataset, ensuring high-confidence ribosome engagement in macrophages. GTFs were generated for both PRICE and RiboTISH ORFs, and candidate gRNAs with cut sites (−4 bp from PAM) intersecting these ORFs were enumerated. GRNAs were filtered to exclude those with MIT specificity score^143^ <50, Ruleset2 cleavage efficiency^144^ <30, or which overlap with annotated CDS regions. Up to 10 gRNAs were selected per nCDS, prioritizing guides targeting the 5’ half of nCDSs >200 bp, the 5’ 75th percentile of nCDSs 100-200 bp, or any position for nCDSs <100 bp to balance expected phenotypic impact with guide availability for short sequences. When >10 suitable guides were available, those with highest specificity scores were selected. For nCDSs with overlapping translated regions (arising from alternative transcripts or frameshifts), gRNAs were assigned to combined “ORF-units” for analysis.

###### Controls

Up to 8 gRNAs targeting non-nCDS regions of each lncRNA were included to distinguish ORF-specific effects from transcript-level regulation. For each targeted uORF, the cognate downstream CDS was also targeted with 6 gRNAs. Positive controls we included 100 essential genes from our previous genome-wide screen^77^ and 236 NFκB pathway regulators (138 from our prior NFκB-GFP sorting screen and 98 from GO terms^145^ GO:0007250, GO:0038061, GO:0004704, and GO:0046696). Protein-coding gene gRNAs were designed using CRISPick^144,146^. The 1,500 non-targeting gRNAs with the lowest measured dropout from our prior screen served as negative controls.

Selected gRNA sequences were 20 nucleotides in length, and the 5’-nucleotide was replaced with “G” to enhance transcription. The final library consisted of 39,748 gRNAs.

###### Lentiviral transduction and selection

The gRNA library was synthesized as an oligonucleotide pool by Twist Bioscience and cloned into a lentiviral vector containing puromycin resistance and BFP reporter cassettes as previously described^77,146^. HEK293T cells were transfected with library plasmid and packaging vectors, and the supernatant was harvested at 72 hours. Three distinct clonal lines of NFkB-GFP^+^, Cas9^+^ immortalized bone marrow-derived macrophages were infected with lentivirus at MOI < 0.3 at day −10. After three days (day −7), lentiviral media was replaced with DMEM +10% FBS media and 10 ug/ml puromycin. After seven days of puromycin selection, each clonal line showed >90% BFP positive cells by flow cytometry, and day 0 samples were collected and stored in freezing media (90% FBS, 10% DMSO).

###### Essentiality screen

For 14 days post-selection, cells were split every ∼3 days to maintain confluency between 60-70% while ensuring >2000X gRNA library coverage. On day 14, cells from each clonal line were collected at >1000X and stored in freezing media.

###### Sorting screen

At day 14, cells from each clonal line were treated with 200 ng/mL Pam_3_CSK_4_. After 24 hours, cells were transferred to FACS buffer to prepare for sorting on a BD FACSAria II as previously described^77^. GFP was excited using a 488-nm laser and detected using a 525/50-nm filter. For each clonal line, the top 20% and bottom 20% of cells were collected as GFP-high and GFP-low samples, with >100X coverage maintained in each sorted sample.

###### Screen sequencing and analysis

Genomic DNA was collected from cell pellets and PCR amplified as previously described^77,147^. Libraries were sequenced, 23-nucleotide single-end, on an Illumina NextSeq 2000 sequencer operated by the UCSC Paleogenomics Lab.

##### CRISPR knockout validation

###### CRISPR KO Experiments

Three gRNAs were chosen to target SYNIR and SEMR ORFs.

SYNIR gRNAs: 1-TGGAGTCTTGATCCCGTCTG, 2-GGGTGAACTGAACTGTAATG, 3-TATGCACAGAAGTATGACTC

SEMR gRNAs: 1-GAAACTAAGATACAGTACCA, 2-AACGTGGCAGGTGATATCAG, 3-ACCAGCCACGCAACCTAACG

And three non-targeting gRNAs were used as negative controls. Complementary ssDNA oligos were ordered from IDT, reconstituted, ligated and cloned into pU6 expression vector. The vector also contained a puromycin resistance cassette and mCherry reporter. Lentivirus was prepared as described above and iBMDM samples were each infected with a single gRNA. Infected cells were puromycin-selected for 7 days when the mCherry population reached >90%. Each gRNA-iBMDM line was treated with 200ng/ml LPS for 6 h and samples were collected for RNA processing.

###### RNA sequencing

Total RNA was isolated from cells using TRIzol reagent (SigmaAldrich, T9424) and Direct-zol RNA Miniprep Plus kit (Zymo Research, R2072), following manufacturer’s instructions. RNA samples were sent to Novogene for 150x150 bp stranded library preparation and were sequenced on a NovaSeq X Plus sequencer.

##### Protein expression and secretion

###### Brip1os translation and secretion

HEK 293T cells were grown in DMEM media amended with 10% fetal bovine serum (FBS) and 1% Penicillin/Streptamycin until confluency and were subsequently transfected with 8ug of Brip1os plasmid as well as the backbone vector as a control. Supernatant and cells were collected 72 hrs post transfection when expression levels were high based on mCherry expression. The supernatant was removed, spun at 300 x g for 5 min. at room temperature (RT) and filtered through a 0.45µm filter. 1 tablet of Protease Inhibitor mix was added to each 10 ml of supernatant. Cells were washed twice with 1x PBS, detached with 0.25% trypsin, collected at 300 x g for 5 min. at RT and lysed in RIPA buffer (50mM Tris-HCL pH 7.0, 1mM EDTA, 0.1% SDS, 0.5% Sodium deoxycholate, 1% NP-40, 150mM NaCl) amended with protease inhibitor (RIPA+). Cell lysates were incubated at 4□ on rotation for 20 min., spun at 14, 000 rpm at 4□ for 10 min. and the soluble portion of the cell lysate was used as a basis for immunoprecipitation (IP). IP of cell lysates and supernatant was performed using V5-Tag (E9H8O) Mouse Monoclonal Antibody Magnetic Bead Conjugate. Beads were initially washed 3x with RIPA buffer. 40ul and 50µl of bead slurry were added to 300µl of cell lysate and 5ml of supernatant and incubated on rotation at 4□ overnight. Beads were collected, washed 5x with RIPA+ buffer. Protein was released from the beads by resuspension in RIPA+ containing 25% LDS and incubation at 70 for 10 min. Beads were collected and 10% of reducing agent was added to each sample before running on SDS-PAGE at 90V for 120 min.and transferred to polyvinylidene difluoride membranes using standard transfer conditions on the Trans-Blot Turbo Transfer System. Membranes were blocked in PBS containing 0.1% (v/v) Tween (PBST) amended with 5% dry milk for 1 hour at RT and probed with primary V5 antibody at a 1:1000 dilution at 4□. Horseradish peroxidase-conjugated goat anti mouse was used as secondary antibody.

##### Single-cell RNA-seq reprocessing

###### Tabula muris/senis scRNASeq reprocessing

To expand the scope of genes available for interrogation in these single-cell atlases, we reprocessed 158,420 cells with reads sequenced with the SMART-Seq2^148^ protocol - a sensitive, full transcript sequencing approach, not restricted to the 3’ end originally published in the Tabula Muris and Tabula Senis projects. For quantifying expression, we used Salmon with (--validateMappings --gcBias --seqBias) against GENCODE annotation vM38. We filtered out cells with fewer than 500 unique genes or less than 50,000 total gene counts, leaving ∼110,000 cells with roughly equal distribution across mouse ages 3-, 18-, and 24-month. Cell annotations, including cell ontology ids, tissue of origin, mouse of origin, etc. were matched from the original publications, and the cell annotation approach is described therein. Gene counts were normalized as ln(1 + counts per million). For clustering, 3000 high-variance genes were selected with the scran package and used to generate 50 principal components separately at distinct mouse ages and globally, yielding four Rdata objects, 3-,18-,24-months, and all ages combined. For each age and for the global set UMAPs were generated with scater with n_neighbors 30 and min_dist 0.5. Full pipeline description and scripts are available in the GitHub repo (Data Availability).

###### Reprocessing of the Immune Dictionary scRNA-seq data

To quantify genes that are missing from the published Immune Dictionary matrices, we reprocessed the raw sequencing data of the Immune Dictionary (GEO GSE202186; BioProject PRJNA834959), an atlas of lymph-node cells from female C57BL/6J mice (11 to 15 weeks old) injected with one of 86 cytokines or PBS (three mice per cytokine, 14 PBS mice), multiplexed with antibody hashtags into 44 10x Genomics 3’ v3 libraries processed in 13 batches. The published matrices were generated with Cell Ranger v3.0 against the 10x mm10-3.0.0 reference (31,053 genes), which does not contain several of the lncRNAs analysed here (AI506816, Synir (D17H6S56E-5), D430020J02Rik and Gm33887). Raw reads were retrieved as SRA files; the cell-barcode read was recovered with fasterq-dump (--include-technical) and identified by its match rate to the 10x v3 barcode whitelist, because read lengths differed between libraries.

Reads were aligned and quantified with STARsolo against GRCm39 with the GENCODE vM39 annotation, emulating Cell Ranger (--soloUMIdedup 1MM_CR --soloUMIfiltering MultiGeneUMI_CR --soloCBmatchWLtype 1MM_multi_Nbase_pseudocounts --clipAdapterType CellRanger4 --outFilterScoreMin 30), with strand-specific counting (--soloStrand Forward), EmptyDrops cell calling (--soloCellFilter EmptyDrops_CR), and exon-level (Gene), exon-plus-intron (GeneFull) and spliced/unspliced (Velocyto) outputs. For a test library, per-gene UMI totals from the exon-level output matched the published matrix (Spearman’s rho = 0.98 across 30,192 genes matched by Ensembl ID). Hashtag counts were taken directly from the hashtag reads (15-bp TotalSeq-A tags, at most one mismatch) for the called cells and demultiplexed with hashsolo; cells assigned to several hashtags or to none were removed, and hashtag calls agreed with the published assignments for 89.6% of cells. Ambient RNA was removed with CellBender remove-background on the exon-level counts (150 epochs; expected cells set to the STARsolo cell count; total droplets included = expected cells + 18,000), and doublets were called per library with scDblFinder and removed. We retained cells with more than 500 detected genes, more than 1,000 UMIs and less than 10% mitochondrial reads, matching the original publication. Cell-type labels were taken from the authors’ annotation (Single Cell Portal SCP2554), matched by library and cell barcode (96.9% of annotated cells recovered), leaving 309,036 cytokine-assigned cells in 25 cell types.

###### Human ERV-derived genes and env-class assignment

Human loci encoding retroviral envelope (env) domains were identified from GENCODE v49]. Two protein sets were scanned: all annotated protein-coding translations longer than 250 amino acids, and open reading frames of at least 750 nt on the sense strand of all lncRNA transcripts. The lncRNA ORFs were identified with ORFfinder (https://www.ncbi.nlm.nih.gov/orffinder/), using the standard genetic code with ATG or alternative start codons. Both sets were annotated with InterProScan 5.76-107.0 using the Pfam 38.0, PANTHER 19.0, CDD 3.21, SUPERFAMILY 1.75, FunFam, NCBIfam, SMART, SFLD and AntiFam member databases at their default thresholds. A sequence was classified as env-containing if it matched at least one of ten signatures specific to retroviral envelope proteins: Pfam PF00429 (ENV polyprotein), PF00517 (retroviral envelope protein, TM/gp41-like) and PF13804 (retro-transcribing virus envelope glycoprotein); PANTHER PTHR10424 (viral envelope protein) and PTHR34313 (ERV-K ENV polyprotein); SUPERFAMILY SSF49830 (ENV polyprotein receptor-binding domain); and the CDD heptad-repeat (HR1-HR2) models cd09851, cd09909, cd09950 and cd09951. Matches were assigned to genes through the GENCODE v49 transcript annotation and pooled per gene, yielding 22 env-containing loci. Seven carry an HGNC ERV designation (ERV3-1, ERVFRD-1, ERVK13-1, ERVMER34-1, ERVV-1, ERVV-2 and ERVW-1); the remaining 15 (12 lncRNA and 3 protein-coding genes) do not.

##### Structure prediction

###### Envelope protein structure homology search

The Alphafold 3 web browser interface was used to predict structures for SEMR B230354K17Rik ncORF8, and D430020J02Rik ncORF1 proteins. For SEMR and D430020J02Rik ncORF1, the predicted signal peptide was not included in the prediction, as this would be cleaved to generate a functional protein. For B230354K17Rik ncORF8, one subdomain was folded with high confidence, while the rest of the protein was predicted at low confidence. The high-confidence structure predictions were searched against the AFDB50 database with FoldSeek.

### QUANTIFICATION AND STATISTICAL ANALYSIS

#### CRISPR screen analysis

Sequenced reads were matched to gRNAs with MAGeCK count with the arguments (--trim-5 1 --sgrna-len 19). In cases where gRNAs targeted overlapping nCDS regions, arising from alternative transcripts or out-of-frame translation, gRNAs were matched to “ORF-units”, e.g. Emsy uORF1:Emsy uORF2. nCDS-loci with less than 4 gRNAs were filtered out and Mageck MLE was used to test for essentiality hits between the day 0 and day 14 samples (--control-sgrna negctrl_ids.txt --norm-method control --permutation-round 10 --remove-outliers --sgrna-efficiency gRNA_scores.tsv), where negctrl_ids.txt is the list of 1,500 negative control gRNAs and gRNA_scores.tsv is the DeepHF^149^ cleavage efficiency prediction for each gRNA.

The gRNA counts from the high and low GFP sorting experiments were normalized with DESeq2 calculated size factors adjustment, and processed in MAUDE as clonal replicates with output reported by the geneLevelStats() function. This resulted in three dataframes, one for each clonal replicate, with z-scores assigned to each nCDS-loci and canonical CDS. The three dataframes were then filtered to a final combined set which removed entries that were not directionally consistent across all three datasets, reducing the number of tested nCDS-loci from 4,891 to 1,434. In this filtered set, replicate aggregated p-values and z-scores were calculated with Stouffer’s method and new padj values were assigned using the Benjamini-Hochberg correction.

#### Differential expression analysis

##### Differential gene expression

Reads were quantified with Salmon128 (--validateMappings --gcBias --seqBias) against GENCODE vM31. Transcript-level abundance estimates were summarized to gene level using tximport with length-scaled TPM counts (countsFromAbundance = “lengthScaledTPM”).

Differential expression was performed with DESeq2138 using default parameters and a two-factor design comparing samples with targeting- and non-targeting-gRNAs.

##### Gene set enrichment analysis

Preranked gene set enrichment analysis (GSEA; Subramanian et al., 2005) was performed separately for each knockout (SYNIR and SEMR) in untreated and LPS-stimulated BMDMs using GSEApy v1.3.1 (Fang et al., 2023). For each comparison, genes were ranked by the Wald statistic from the DESeq2 analysis of knockout versus control cells; all genes with a non-missing test statistic were included (approximately 20,000–21,000 genes per comparison). Gene sets were selected a priori, based on the hypothesized roles of the targeted loci in innate immune and NFκB signaling, from the Reactome, BioCarta and WikiPathways collections of MSigDB (mouse, v2024.1–v2025.1), and covered innate immune and NFκB signaling, cytokine and interferon responses, inflammasome and complement pathways, antigen presentation, cell death and metabolism. Gene sets with fewer than 5 or more than 500 genes present in the ranked list were excluded, leaving 36 gene sets (27 Reactome, 7 BioCarta, 2 WikiPathways) for testing. Enrichment scores were calculated with the weighted running-sum statistic, and normalized enrichment scores (NES) were obtained by dividing each enrichment score by the mean of the permutation-derived scores of the same sign. Significance was estimated from 1,000 gene-label permutations, and false discovery rate (FDR) q-values were computed by the standard GSEA procedure, in which each observed NES is compared with the null NES distribution pooled across all tested gene sets, separately for positive and negative scores.

Gene sets with FDR < 0.10 in either condition are shown, with non-significant results (FDR ≥ 0.10) in the other condition indicated in grey. Gene sets with fewer than 10 genes, and gene sets related to translation, nonsense-mediated decay and DNA double-strand break repair, were included in the analysis but omitted from the figures.

##### Differential expression of ERV genes in immune cytokine dictionaries

Mouse. For each cell type, CellBender-corrected exon-level counts were summed into one pseudobulk profile per mouse. PBS mice were split across all libraries of their processing batch, so their cells were pooled across libraries to give one sample per animal (one library contained two PBS mice, which were combined into one sample). Samples with fewer than 10 cells were removed, and cytokines with at least two remaining mice were tested; samples of the other cytokines were removed before fitting. This left two or three mice per tested cytokine, 81 to 86 tested cytokines and 10 to 13 PBS samples per cell type. Differential expression against PBS was tested with edgeR (v4.8.2) quasi-likelihood F-tests. Genes were filtered with filterByExpr, retaining 14 genes regardless of^110^ expression: the ERV-derived genes AI506816, Synir, Brip1os, B230354K17Rik, D430020J02Rik, Fv1, 2410002F23Rik and Gm33887, the syncytin genes Syna and Synb, and the interferon-stimulated genes Irf7, Isg15, Stat1 and Mx1. Libraries were TMM-normalized, and models with the design ∼ batch + cytokine (processing batch as a covariate; PBS as the reference level) were fitted with glmQLFit using its default settings. Each cytokine was compared with PBS by a quasi-likelihood F-test. Syna and Synb had only 4 and 9 counts across pseudobulks and were not tested. For the dot plots (Fig. 4F and 5B), P values were adjusted with the Benjamini-Hochberg procedure within each cell type, over all gene x cytokine tests of the eight tested ERV-derived genes, and FDR < 0.05 was considered significant.

Human. Human data were from the human cytokine dictionary^93^ (Parse Biosciences; 9,697,974 peripheral blood mononuclear cells from 12 donors stimulated in vitro with 90 cytokines or PBS; genes quantified against Ensembl release 109), downloaded as the released h5ad file (https://www.parsebiosciences.com/datasets/10-million-human-pbmcs-in-a-single-experiment/). Raw counts were summed into pseudobulks per donor, cytokine and cell type, using the 18 cell-type labels and the cytokine labels of the released data. Pseudobulks with at least 10 cells were kept, and cytokines with at least two donors in a cell type were tested. Differential expression against PBS was tested with edgeR quasi-likelihood F-tests with the design ∼ donor + cytokine (PBS as the reference level). Genes were filtered with filterByExpr, retaining the 142 ERV-derived genes present in the data (see below) regardless of expression; libraries were TMM-normalized; and models were fitted with glmQLFit using its default settings. Each cytokine was compared with PBS by a quasi-likelihood F-test, and P values were adjusted with the Benjamini-Hochberg procedure within each cell type over all ERV-derived gene x cytokine tests (FDR < 0.05). The interferon-stimulated genes IRF7, ISG15, STAT1 and MX1 were tested from the same fits.

---

**Supplemental Figure 1.**
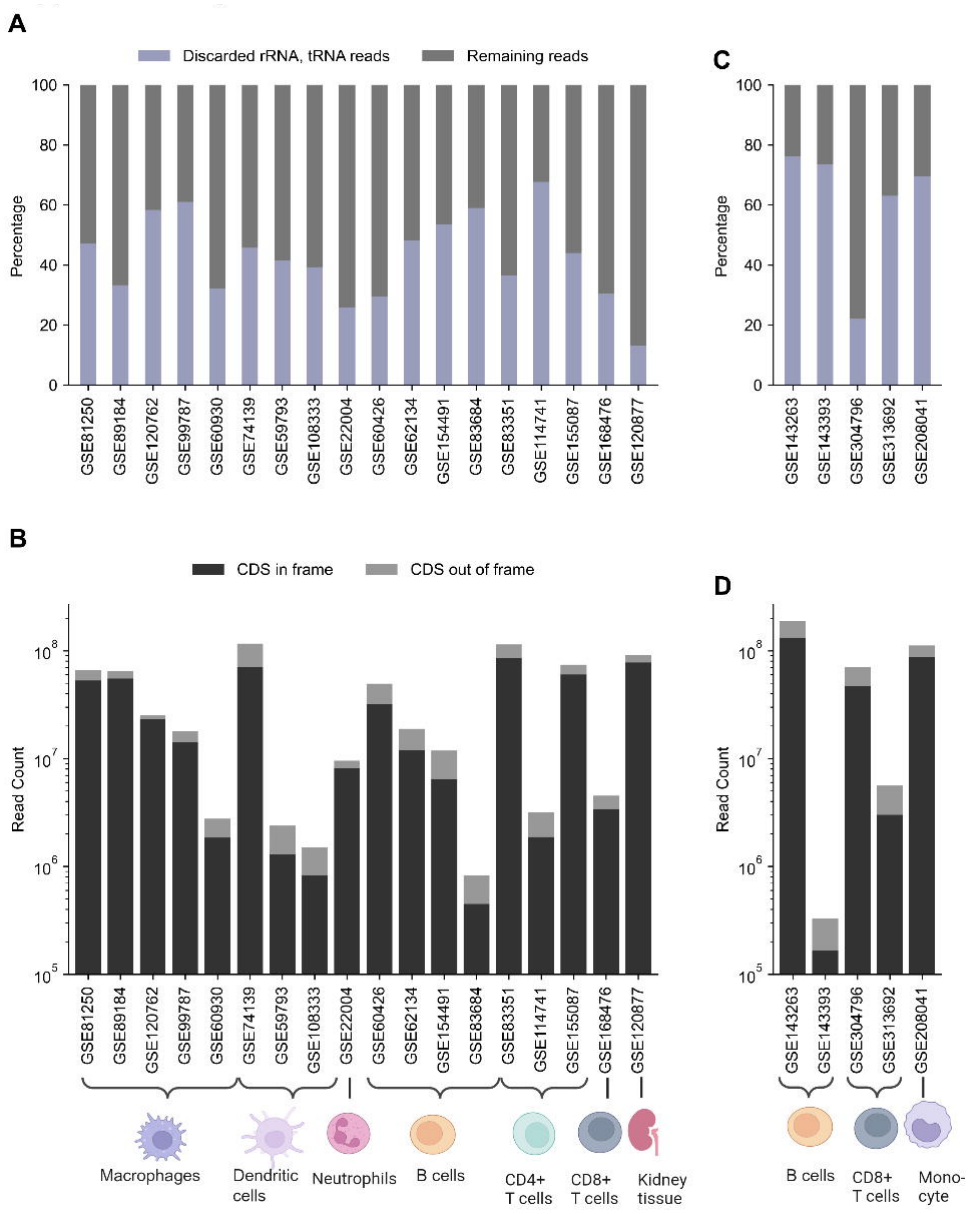
**Quality metrics of mapped Ribo-seq reads by dataset**. (A,C) QC on Ribo-seq showing proportion of reads mapping to rRNA and tRNA (discarded) versus reads that can be used to call translation events for (A) mouse and (C) human (B,D) Counts for Ribo-seq reads after filtering, with the number of reads mapping in-frame to known CDS p-sites vs out-of-frame in the CDS (B) mouse and (D) human CDS: coding sequence

**Supplemental Figure 2.**
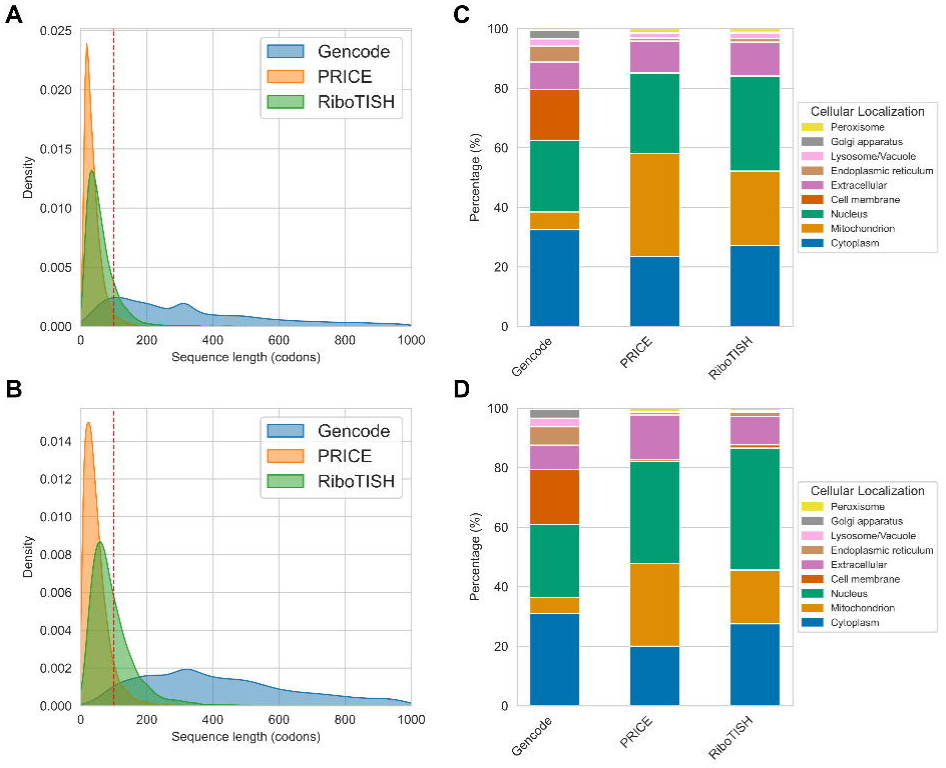
L**e**ngth **and localization predictions of non-canonical CDSs.** (A,B) NCDS lengths from annotated GENCODE vM31 coding sequences (mouse) and GENCODE v49 (human) and predicted nCDSs from PRICE and RiboTISH for (A) mouse and (B) human. Dashed line at 100 codons references heuristic length-based coding sequence cutoff. (C, D) DeepLoc predictions of localization compartments of the nCDS-encoded proteins of annotated or predicted coding sequences for (C) mouse and (D) human.

**Supplemental Figure 3.**
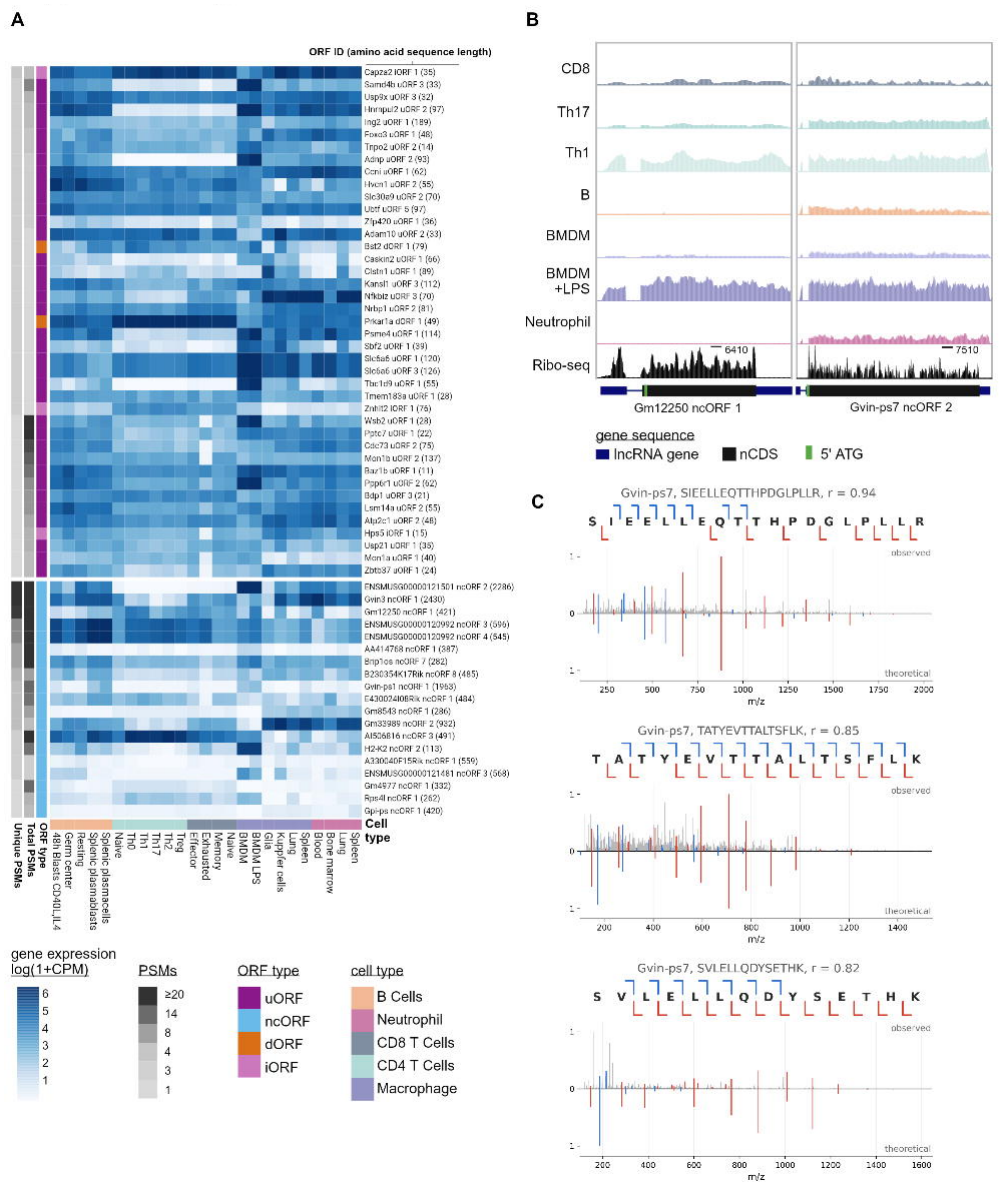
P**r**oteomics **supports translation of select noncanonical loci** .(A) RNA expression Heatmap of nCDS with either a minimum of 5 total peptide spectra matches (PSMs) or 2 unique PSMs. Gene expression is uniformly processed and batch corrected with Limma to calculate comparable counts per million (CPM). Final expression for a given cell type is the mean of experimental replicates. (B) Uniquely mapped bulk RNA and Ribo-seq at ENSMUSG00000121501 (or Gvin-ps7) and Gm12250, interferon induced pseudogenes with strong evidence of transcription and translation. (C) Peptide spectra that map uniquely to Gvin-ps7, nowhere else in the genome, passed FDR filters, and scored “good” according to the heuristic scorer (Methods).

**Supplemental Figure 4.**
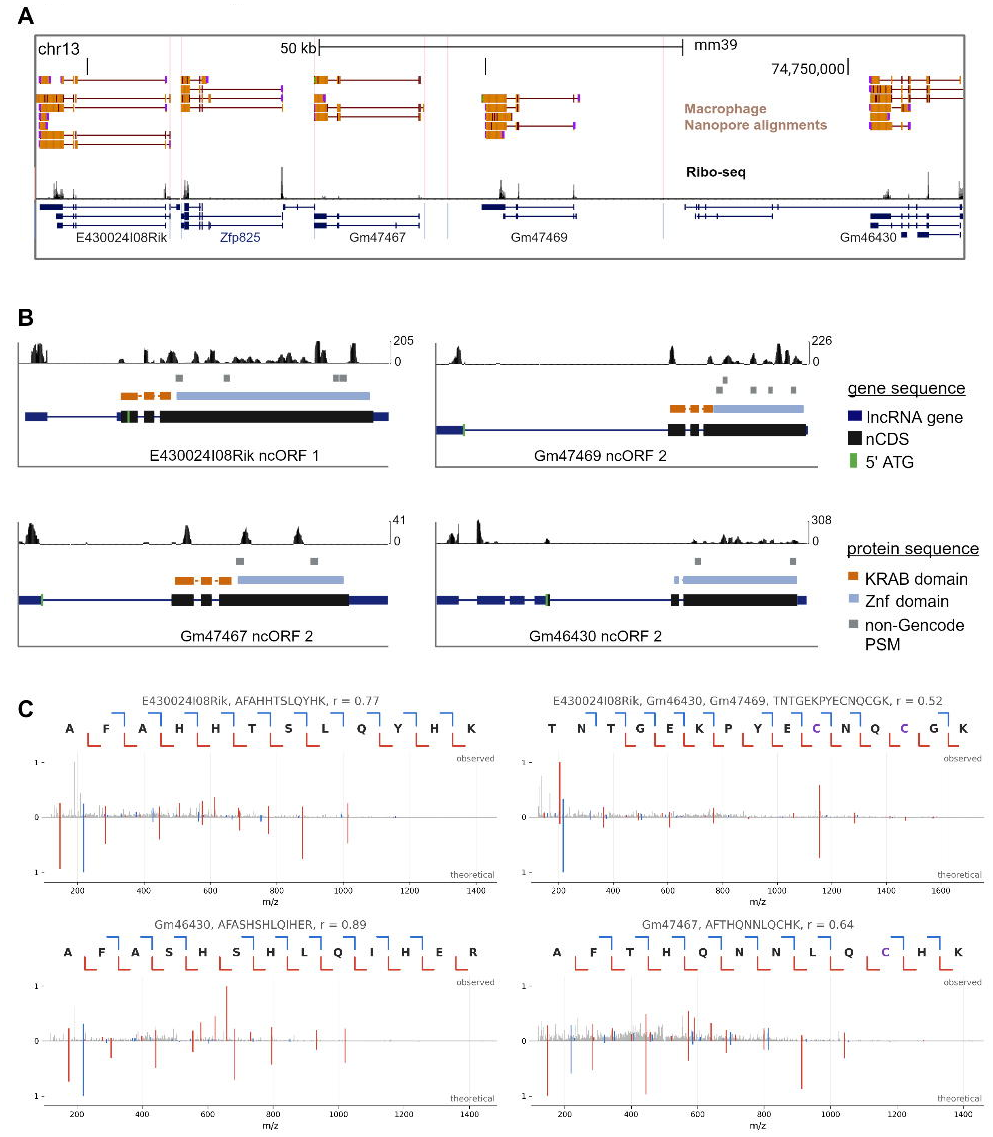
U**n**annotated **Zinc finger protein cluster on chromosome 13**. (A) Uniquely mapped Nanopore direct RNA reads and Ribo-seq reads at a cluster of neighboring zinc finger proteins. Zfp825 is annotated as a protein coding gene while the others are long noncoding RNAs. (B) Ribo-seq reads and predicted KRAB and zinc finger domains at four long noncoding RNAs neighboring Zfp825. (C) Peptide spectra that passed FDR filters, do not map to a known GENCODE protein, and scored “excellent” according to the heuristic scorer (Methods).

**Supplemental Figure 5.**
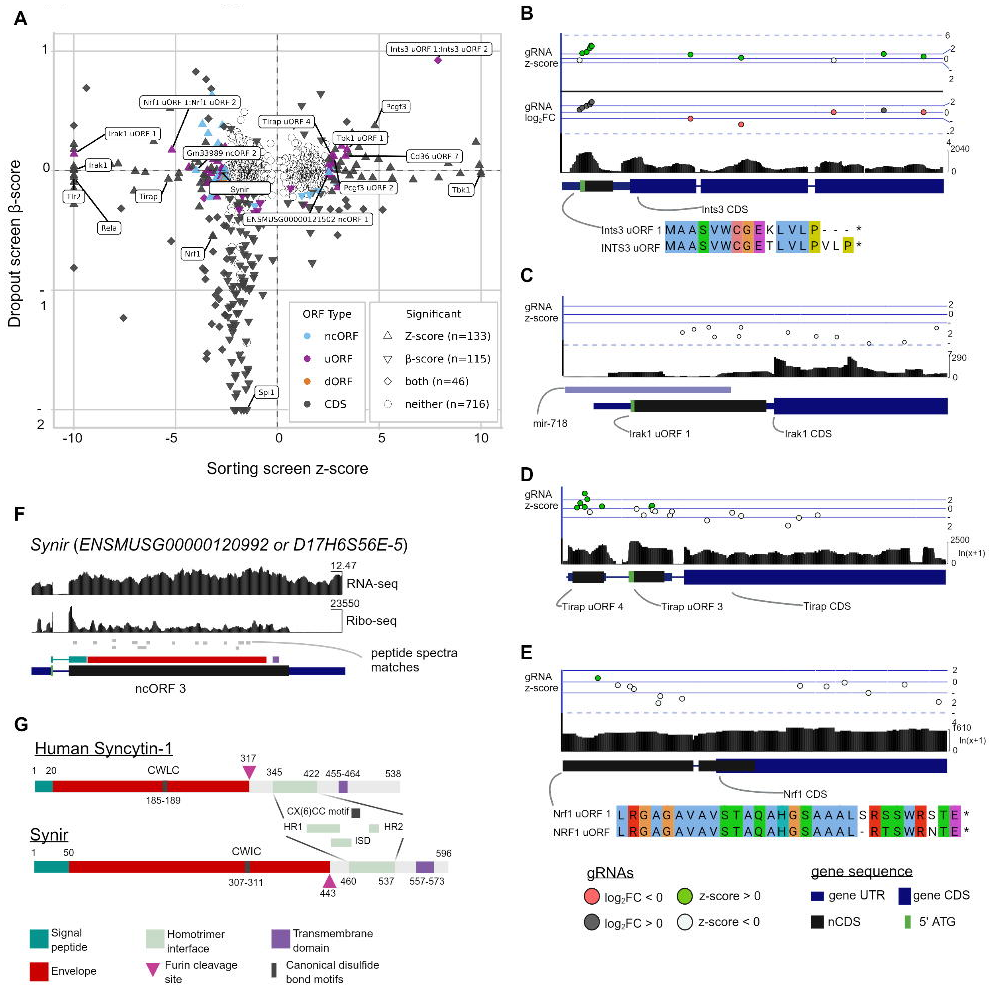
C**o**mbined **screen results** (A) The results of targeting nCDS across both screens. The majority of significant hits are screen-specific. To enhance visualization, beta scores < −2 were capped at −2 and z-scores beyond +/-10 were capped at −10 and 10. Full values are available in the associated table. (B-E) NCDS loci illustrating gRNA combined z-score and dropout (Stouffer’s mean from triplicates), Ribo-seq support, ORF structure and conservation with human ORF where applicable. Mouse Ribo-seq is combined from all studies (Figure 1). Ranges are raw signal counts. (B) Ints3 uORF was a significant hit in both screens (dropout: Wald p-value < 0.02, padj. < 5E-4 ; sorting: mean z-score > 7.8, padj < 9.3E-14) as a growth suppressor and negative NFkB regulator, while Ints3 was not a significant hit in either screen. (C) Irak1 uORF 1 was the most significant novel ORF regulator of NFkB (z-score < −9.9, padj < 6.9E-299), but the gRNAs are highly overlapping with microRNA 718, confounding a direct link for the ORF. (D) Tirap uORF 4 is grouped with negative regulators of NFkB (mean z-score > 2.6, padj < 3.2E-2) while Tirap primary coding sequence (mean z-score < −5.3, padj < 1.8E-6) is grouped with positive regulators. Tirap uORF 3 is highly translated, but does not show a consistent effect across replicates. Signal track is log-transformed to improve visualization. (E) Nrf1 uORF 1 was ranked as a more significant regulator of the pathway (z-score = −5.2 padj < 4E-06) than Nrf1 coding sequence (z-score = −3.2, padj < 0.01). For (D,E) signal track is log-transformed to improve visualization. (F) Synir genomic locus with Ribo-seq, BMDM RNA-seq, peptide spectra matches, and predicted protein domains. (G) Shared domains and features between SYNIR protein and human Syncytin-1. Signal peptide predicted by Signal6P and DeepTMHMM. Transmembrane domain predicted with DeepTMHMM. Furin cleavage site predicted with ProP. Other domains predicted by Interproscan. ISD: immunosuppressive domain; HR1: heptad repeat 1 domain; HR2: heptad repeat 2 domain; LPS: lipopolysaccharide

**Supplemental Figure 6.**
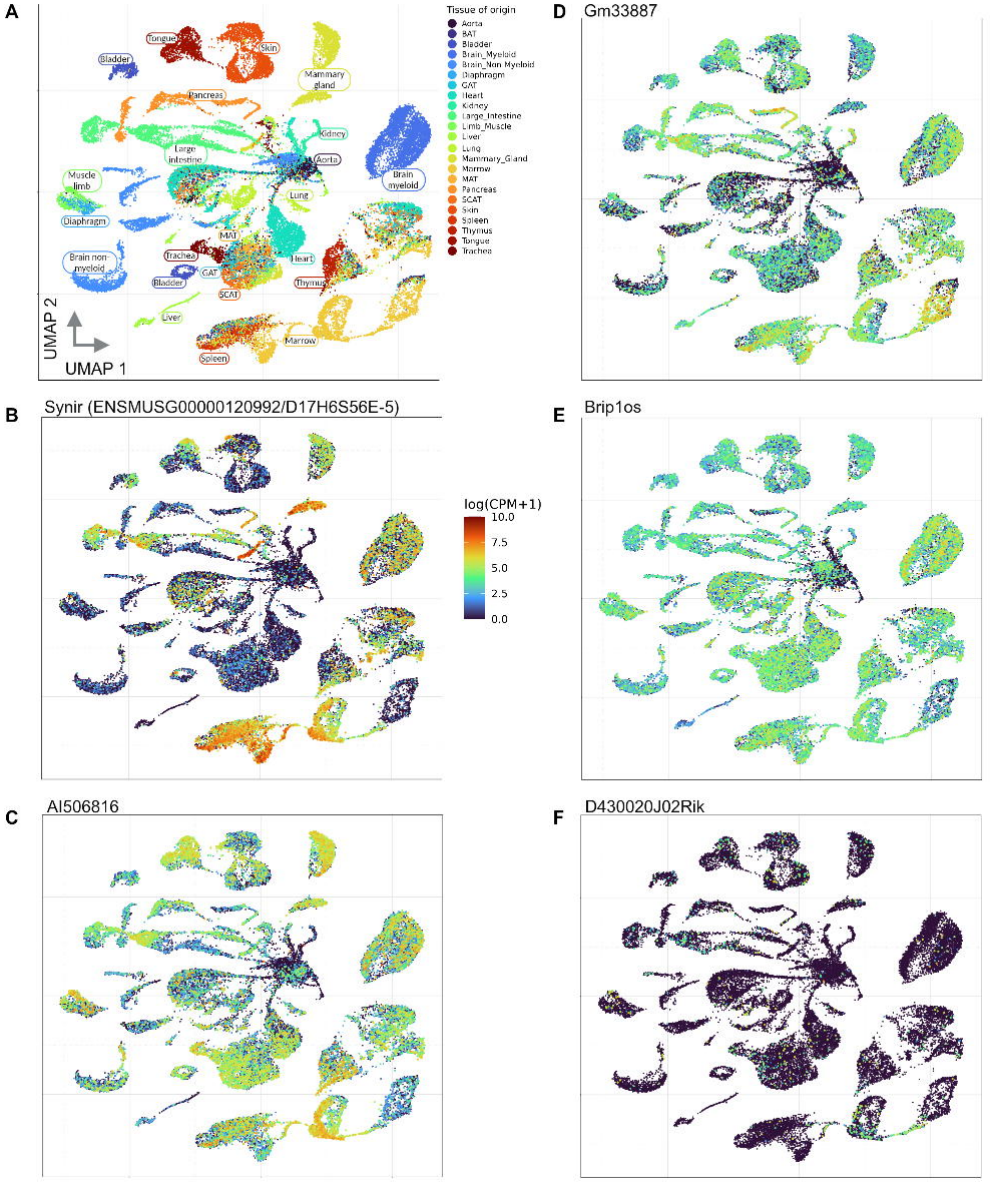
T**a**bula **Muris/Senis scRNA-seq reanalysis including lncRNAs highlights retroviral-envelope-like gene expression across tissues.** (A) Reference UMAP of single-cell RNA-seq colored by tissue of origin. (B-F) (B) Synir (*ENSMUSG00000120992*/*D17H6S56E-5*), (C) *AI506816*, (D) *Gm33887*, (E) *Brip1os*, and (F) *D430020J02Rik*, which are predicted to be derived from retroviral envelope genes.

**Supplemental Figure 7.**
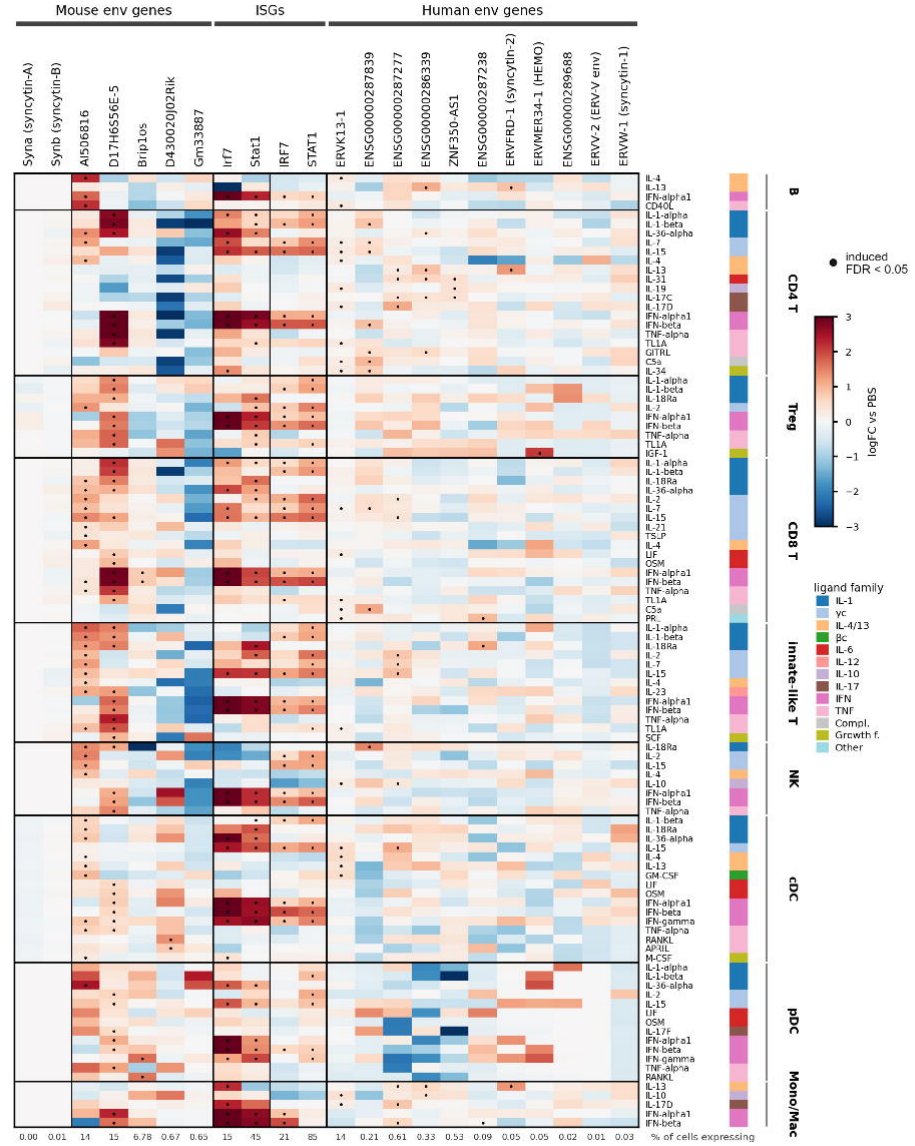
Cytokine responses of envelope-class ERV-derived genes in mouse and human immune cells. Heatmap of cytokine-induced expression changes for mouse env genes, interferon-stimulated gene (ISG) anchors and human env-class ERV-derived genes (columns) across 104 lineage-ligand stimuli (rows). Mouse data are from the Immune Dictionary (lymph node after in vivo cytokine injection; one pseudobulk per mouse, three mice per cytokine) and human data from the human cytokine dictionary (peripheral blood mononuclear cells stimulated in vitro; one pseudobulk per donor, 12 donors); in both, expression was compared with PBS using edgeR quasi-likelihood F-tests. Colour shows logFC versus PBS, averaged over the cell types that make up each lineage and clipped at ±3. Dots mark induction (FDR < 0.05 and logFC > 0 in at least one constituent cell type), with Benjamini-Hochberg correction across all gene x cell type x ligand tests of each species’ panel (mouse: 8 tested ERV-derived genes and 4 ISG anchors; human: 99 env- or gag-class genes and 4 ISG anchors). Mouse env genes: syncytin-A (Syna), syncytin-B (Synb), AI506816, D17H6S56E-5, Brip1os, D430020J02Rik and Gm33887; Syna and Synb (fewer than 100 counts in total) are shown with their logFC but were not tested. ISG anchors: mouse Irf7 and Stat1, human IRF7 and STAT1. Human env genes: the 11 genes with an env domain in the released data, ordered by number of inductions; ERVV-1 is absent because it was filtered from the released data (detected in fewer than 200 cells). Rows include every stimulus that induces a mouse ERV-derived gene, induces at least two human env-class genes, or induces ERVW-1, ERVFRD-1, ERVV-2 or ERVMER34-1; they are grouped by lineage (brackets, right) and marked by ligand family (colour strip). Ligand names follow the human dataset (IL-18 appears as IL-18Ra). Numbers below the columns give the percentage of cells expressing each gene (mouse, 576,463 lymph-node cells; human, 9,697,974 PBMCs).

**Supplemental Figure 8.**
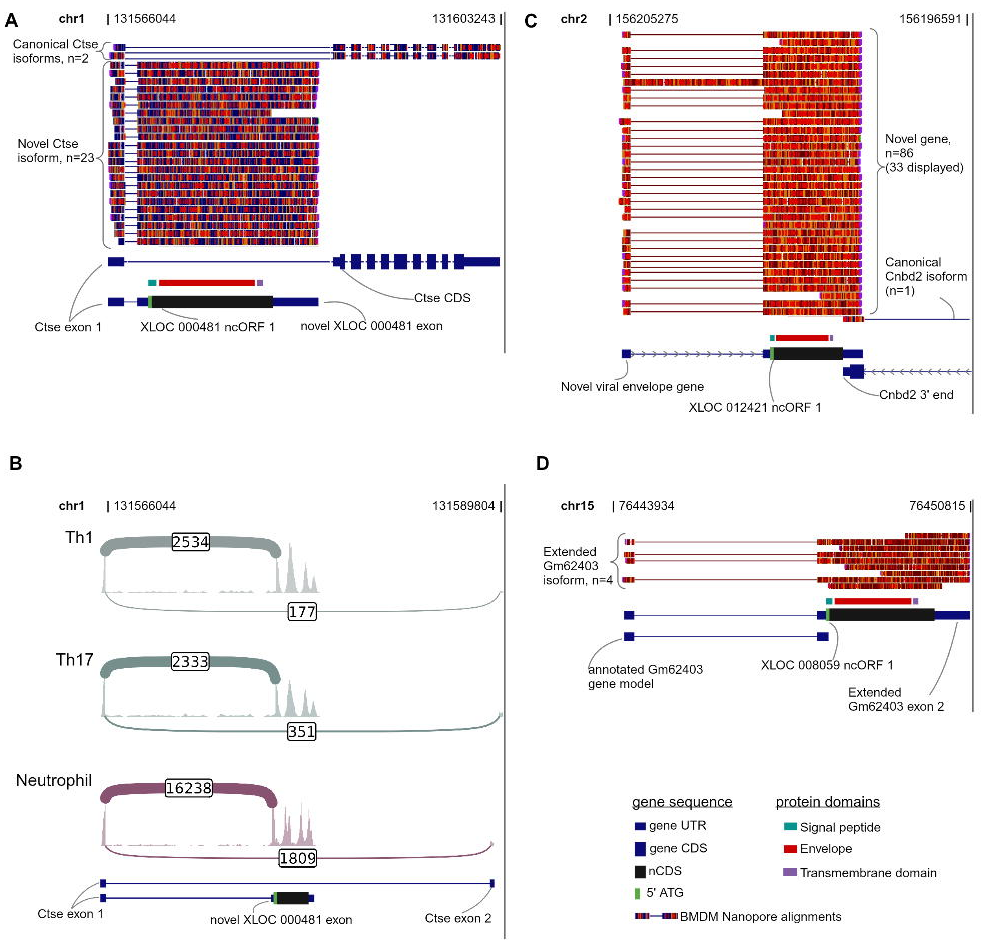
N**a**nopore **long read support for unannotated retroviral loci and isoforms.** (A) Nanopore long reads shows a dominant second exon in Ctse that is highly expressed relative to the canonical isoform. (B) Sashimi plots showing short read RNA support for expression of the novel *Ctse* isoform. (C) Nanopore long read support for an unannotated retroviral gene antisense to *Cnbd2* (D) Nanopore long read support for an extended second exon encoding a retroviral envelope protein in lncRNA *Gm62403*.

**Supplemental Figure 9.** I**n**teractive **and customizable public Genome Browser session for further inquiry.**

(B) Example of the Leukocyte ORF Genome Browser session at the Brip1os locus. The session includes (from top to bottom) guide RNA effect sizes, mouse and human Ribo-seq, protein domain predictions, peptide spectra matches, nCDS predictions, GENCODE gene annotations, bulk RNA-seq and cross-species conservation

(C) Guide RNA effect sizes for both dropout and sorting screen can be displayed as means or per replicate.

(D) Ribo-seq tracks can be set to show mouse (aggregate leukocyte) and human (THP1) signal tracks from elongation inhibition experiments (cycloheximide) as well as signal from translation initiation inhibition experiments (lactimidomycin or harringtonine).

(E) Proteomics tracks include domain predictions from SignalP (secreted peptides), DeepTMHMM (secreted peptides and transmembrane proteins), Interproscan (protein domain database aggregator) and peptide spectra matches from comparison

(F) NCDS prediction tracks include merged PRICE and RiboTISH calls as well as per-tool tracks and configurable display of transcriptome sources.

